# RAD51AP1 is a versatile RAD51 modulator

**DOI:** 10.1101/2025.05.24.655746

**Authors:** Lucas Kuhlen, Bilge Argunhan, Pengtao Liang, Janet Zhong, Laura Masino, Xiaodong Zhang

**Author notes:** These authors contributed equally to this work (Lucas Kuhlen, Bilge Argunhan).

## Abstract

RAD51AP1 is an emergent key factor in homologous recombination (HR), the major pathway for accurate repair of DNA double-strand breaks, and in alternative lengthening of telomeres (ALT). Depletion of RAD51AP1 diminishes HR and overexpression is common in cancer, where it is associated with malignancy. Here, we show that RAD51AP1 serves as a versatile modulator of the RAD51 recombinase, the central player in HR. Through a combination of biochemistry and structural biology, we reveal that RAD51AP1 possesses at least three RAD51-binding sites that facilitate its binding across two adjacent RAD51 molecules. We uncover a novel RAD51 binding mode and characterise a hitherto overlooked role for RAD51AP1 in stabilising RAD51-ssDNA filaments and promoting strand exchange. Further, we resolved structures of RAD51-ssDNA filaments in the presence of Mg^2+^-ATP and Mg^2+^-ADP, revealing conformational changes upon ATP hydrolysis and explaining how ADP reduces RAD51-DNA binding. Our findings provide mechanistic insights into RAD51 recombinase and RAD51AP1.

**Highlights:**

1. Structures of RAD51 filaments in the presence of Mg^2+^-ATP and Mg^2+^-ADP reveal how ATP hydrolysis reduces DNA binding
2. ATP hydrolysis does not induce RAD51 filament contraction, instead it induces filament relaxation
3. RAD51AP1 uses three sites to bind across two RAD51 monomers
4. RAD51AP1 uses a unique binding mode to stabilise the RAD51 N-terminal domain and protomer interface of RAD51
5. RAD51AP1 binding induces conformational changes that promote RAD51 DNA association, oligomerisation, filament nucleation, filament stabilisation and strand exchange

## Introduction

The human genome is constantly being damaged and repaired. DNA double-strand breaks (DSBs) constitute the most cytotoxic form of DNA damage, and as a result, their faithful repair is critical for the maintenance of genome stability. Homologous recombination (HR) is the major pathway for high-fidelity repair of DSBs (Symington and Gautier, 2011; Wright et al., 2018). Disruption of HR leads to genomic instability, which is a strong driver of tumorigenesis (Prakash et al., 2015). Furthermore, HR-directed repair is one of the major pathways to resolve stalled or damaged replication forks and for alternative lengthening of telomeres (ALT) (Barroso-Gonzalez et al., 2020; O’Sullivan and Greenberg, 2024; Tye et al., 2021).

The central protein in HR is the RAD51 recombinase. RAD51 forms a nucleoprotein filament with single-stranded DNA (ssDNA) generated from enzymatic end resection at the DSB site (Cejka and Symington, 2021; Gnugge and Symington, 2021). The RAD51-ssDNA filament performs the recombinase function through searching and identifying homologous sequences in intact donor double-stranded DNA (dsDNA), invading the dsDNA to allow the pairing of the ssDNA with the complementary strand from donor DNA, and using it as a template for DNA synthesis, resulting in repair (Haber, 2018; Sun et al., 2020). However, unlike its bacterial orthologue RecA, the intrinsic recombinase activity of RAD51 is low, and eukaryotic HR requires a multitude of auxiliary factors to promote RAD51 activities for proficient DSB repair (Carver and Zhang, 2020). Furthermore, erroneous HR can lead to unregulated strand invasion and chromosome instability (Klein, 2008; Piazza et al., 2017), and chromosomal accumulation of RAD51 can lead to cell toxicity (Carver et al., 2024; Klein, 2008). RAD51’s activity thus needs to be tightly regulated. One emergent RAD51 modulator is RAD51AP1 (RAD51 Associated Protein 1), a protein that physically binds to RAD51 (Kovalenko et al., 1997) and promotes its activities in HR, thus contributing to the maintenance of genome stability (Modesti et al., 2007; Wiese et al., 2007). Importantly, RAD51AP1 is overexpressed in many types of cancer, including breast and ovarian cancer, and this overexpression is associated with a poor prognosis (Pires et al., 2017). RAD51AP1 deletion improves survival in mouse models of breast cancer and its expression is associated with cancer stem cell self-renewal, which is highly dependent on functional DNA repair (Bridges et al., 2020), confirming its important roles in HR and cancer development. More recently, RAD51AP1 has been shown to play key roles in R-loop formation and ALT (Kaminski et al., 2022; Ouyang et al., 2021; Yadav et al., 2022).

Sequence and secondary structure analysis as well as biochemical investigation (Modesti et al., 2007) suggest that RAD51AP1 is a largely intrinsically disordered protein (**Fig. 1A, Fig. S1A**). Earlier biochemical and genetic studies identified a RAD51 binding site and two DNA/RNA binding sites (**Fig. 1A**), which are all required for full function in cells (Kaminski et al., 2022; Ouyang et al., 2021; Wiese et al., 2007). In HR, it is thought that RAD51AP1 engages with RAD51 through its C-terminal RAD51 binding site (Kovalenko et al., 2006; Wiese et al., 2007) and then captures dsDNA with its DNA binding sites to form a molecular bridge between the filament and the strand exchange donor (Modesti et al., 2007; Wiese et al., 2007). In ALT, RAD51AP1 has been proposed to bind to telomere-specific RNA (TERRA) and promote R-loop formation, which subsequently promotes D-loop formation and HR-directed repair (Barroso-Gonzalez et al., 2020; Ouyang et al., 2021; Yadav et al., 2022) ; it is unclear if RAD51 binding activities also play a role here. RAD51AP1 has also been shown to bind to nucleosomes, supporting the notion that it can promote the pairing of a DSB site with the sister chromatid in cells (Pires et al., 2021).

**Figure 1.**
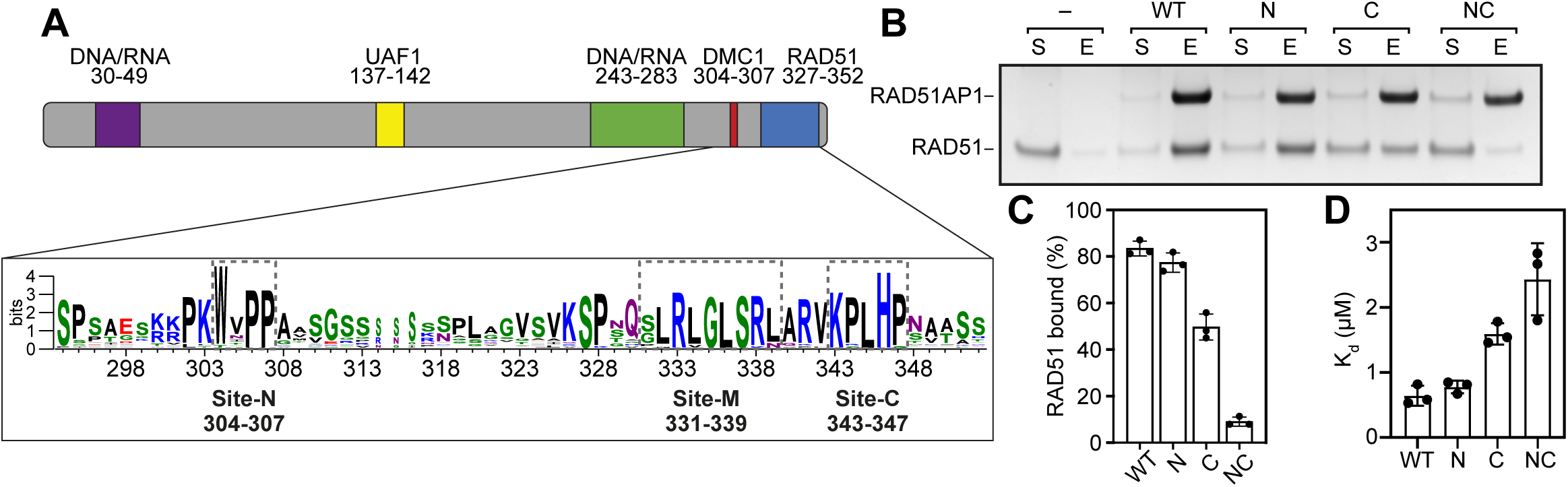
RAD51AP1 binds to RAD51 via two distinct binding sites. **(A)** Schematic of RAD51AP1 showing regions of interest. **(B)** Pull-down experiment of full-length RAD51AP1 (6xHis-tagged wild type or mutants) and RAD51. **(C)** Quantification of **(B)**. Averages are shown with error bars depicting standard deviation (n = 3). **(D)** Kd values for the binding of full-length RAD51AP1 and RAD51, determined by bio-layer interferometry. Averages are shown with error bars depicting standard deviation (n = 3).

Despite its importance in HR and other processes that involve RAD51, the exact molecular mechanism of RAD51AP1 in these activities is unknown. Importantly, thus far, due to the instability of RAD51-DNA filaments in the presence of Mg^2+^ and ATP, current structures of RAD51-DNA filaments have been resolved either by replacing Mg^2+^ with Ca^2+^, which curbs ATP hydrolysis (Bugreev and Mazin, 2004), or using the non-hydrolysable ATP analogue AMP-PNP (Robertson et al., 2009) or the hydrolysis deficient RAD51-K133R mutant (Chi et al., 2006; Morrison et al., 1999) (Appleby et al., 2023b; Xu et al., 2017). Single-molecule analysis has suggested that ADP-bound protomers dissociate from the filament (van Mameren et al., 2009), providing a feasible explanation for why restricting ATP hydrolysis may lead to enhanced filament stability.

Here, using cryo electron microscopy, we resolved structures of RAD51-ssDNA filaments in complex with Mg^2+^-ATP and Mg^2+^-ADP, revealing conformational changes upon ATP hydrolysis and the structural basis for their distinct DNA binding properties. Furthermore, we obtained structures of RAD51AP1 in complex with RAD51-ssDNA filaments, uncovering the mechanism of RAD51AP1 in RAD51 filament modulation. Combined with biochemical studies and mutagenesis, we show that RAD51AP1 possesses three RAD51-binding sites that facilitate its binding across two adjacent RAD51 molecules. We uncover a novel RAD51 binding mode, as well as characterise a hitherto overlooked role for RAD51AP1 in stabilising RAD51-ssDNA filaments and re-ordering structural elements in RAD51 that promote strand exchange. Our studies reveal that RAD51AP1 acts as a RAD51 modulator via strengthening protomer-protomer interactions, promoting RAD51 oligomerisation and re-ordering its DNA binding sites, thereby facilitating filament nucleation, stabilisation and strand exchange.

## Results

### RAD51AP1 binds to RAD51 via distinct sites

Previous biochemical analyses identified a RAD51 binding site near the extreme C-terminus of RAD51AP1, with the H346A mutation severely impairing RAD51 binding (Wiese et al., 2007). Upon examination of the RAD51AP1 sequence, we noticed a stretch of conserved residues upstream of this binding site (**Fig. 1A, S1B**). Interestingly, these residues were previously shown to comprise the binding site for DMC1, the meiosis-specific RecA-family recombinase in eukaryotes (Dunlop et al., 2011). We refer to this DMC1 binding site as Site-N, and the known RAD51 binding site as Site-C for their relative positions in the sequence. Given the structural and sequence similarity between RAD51 and DMC1, we hypothesised that Site-N may also be involved in RAD51 binding. To test this, we mutated five conserved residues (K300,K303,W304,P306,P307) in Site-N to alanine. We also included the known mutant (RAD51AP1-H346A) that disrupts RAD51 binding in our analysis as a Site-C mutant, and designed a RAD51AP1 variant in which both sites are mutated (referred to as NC). Along with RAD51, these RAD51AP1 variants were purified to homogeneity (**Fig. S1C**).

As expected, RAD51 can be pulled down by RAD51AP1 and the Site-C mutant showed a reduction in binding (**Fig. 1B-C**). Compared with wild-type RAD51AP1, the Site-N mutant did not show an appreciable reduction in RAD51 binding. Interestingly, when both binding sites were mutated, the binding was almost abolished (**Fig. 1B-C**). To quantify these interactions, biolayer interferometry (BLI) experiments were conducted and the results are consistent with those of pull-down experiments (**Fig. 1D, S1D-E**). Wild-type protein has a dissociation constant of ∼0.6 μM, which is comparable to the Site-N mutant, whereas the Site-C mutant decreased the binding affinity by ∼2.5 fold, and the Site-NC double mutant showed a ∼4-fold reduction in affinity (**Fig. 1C**). These results indicate that RAD51AP1 binds to RAD51 via at least two distinct sites, and these two sites synergistically enhance the binding, although Site-C is the major binding site.

### RAD51AP1 promotes RAD51 filament nucleation, stabilises filaments and promotes strand exchange

Given that RAD51AP1 is known to promote HR via potentiation of RAD51, we next tested if it can stimulate RAD51 activities including filament nucleation, stabilisation, and strand exchange. To directly assess the effects of RAD51AP1 on RAD51-ssDNA filaments, we used negative stain electron microscopy to visualize and quantify the RAD51 filaments formed in the presence of Mg^2+^-ATP (**Figs. 2A-C**). Because the addition of RAD51AP1 to filaments caused aggregation, we employed a C-terminal fragment that contains both binding sites (C59, similar to C60 used in (Dunlop et al., 2011)) that did not cause aggregation and binds to RAD51 in a similar fashion as the full length protein (**Fig 1B**, **2D**). Similar to previous studies of other RAD51 modulators such as BRCA2 (Shahid et al., 2014) and fission yeast Swi5-Sfr1 (Liang et al., 2023), addition of RAD51AP1-C59 led to an increase in the number of filaments observed and shifted the distribution of filament lengths towards longer filaments (**Figs. 2B-C**), suggesting it increases filament nucleation and stability. Interestingly, mutation of either Site-N or Site-C caused a reduction in filament number (**Fig. 2B**), but only mutation of Site-C abolished the increase in filament length conferred by RAD51AP1-C59 (**Fig. 2C**).

**Figure 2.**
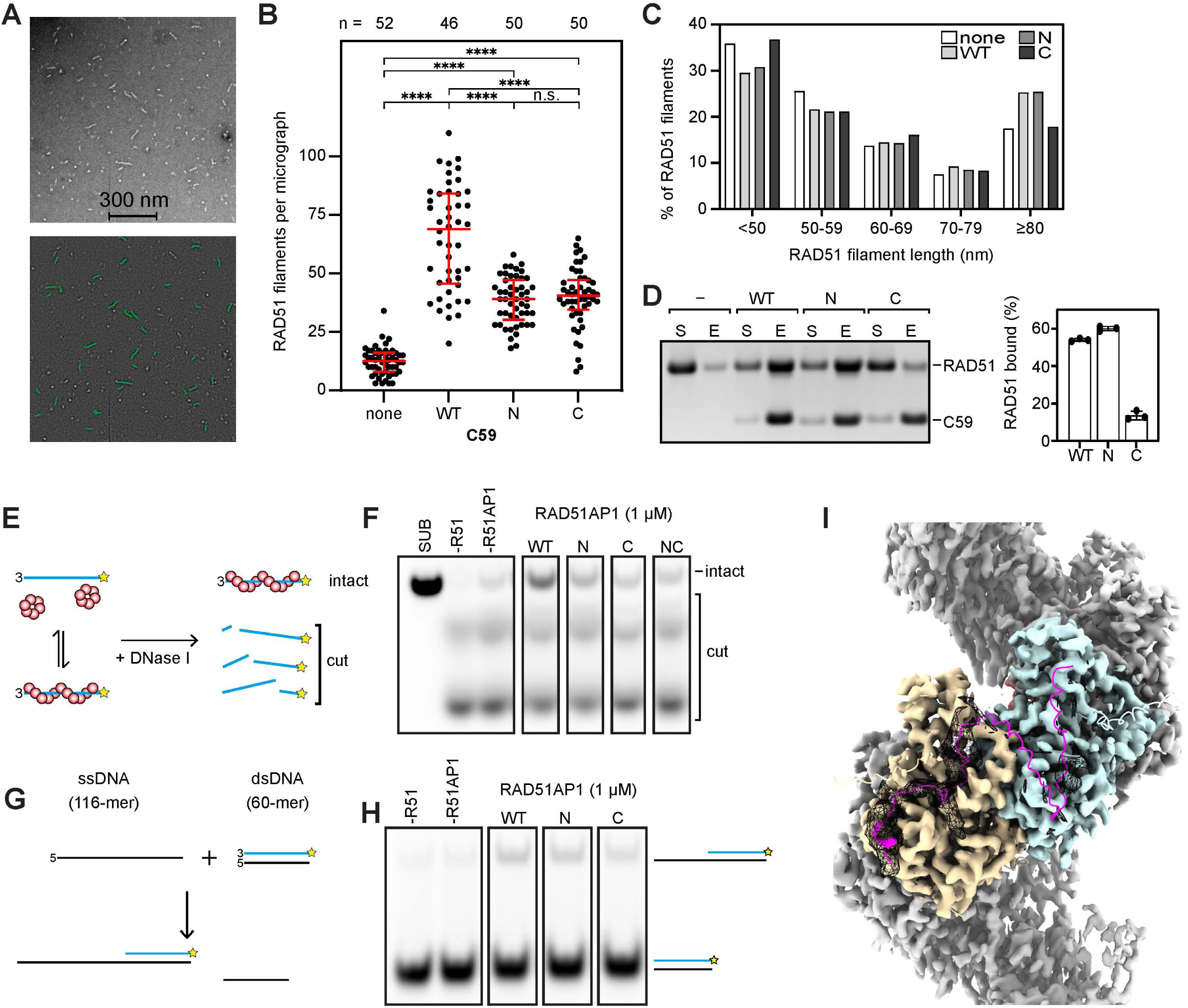
RAD51AP1 promotes filament formation, stability, and strand exchange. **(A)** Negative-stain electron micrographs of RAD51 filaments. RAD51 filaments were prepared either with RAD51 alone (none) or with a variant of C59 (WT, N, C) added. Top panel shows the captured image, bottom panel shows the image after automated processing to measure filament length and number. **(B)** Quantification of RAD51 filaments per micrograph. Red bar represents the median with lower and upper quartiles. **(C)** Quantification of RAD51 filament length. Filaments were categorised into the indicated lengths and expressed as a percentage of the total. none, n = 642; WT, n = 2997; N, n = 1918; C, n = 1993. **(D)** Pull-down experiment of RAD51AP1-C59 (6xHis-tagged wild type or mutants) and RAD51. For quantification, averages are shown with error bars depicting standard deviation (n = 3). **(E)** Schematic of nuclease protection assay to assess RAD51 filament stability. Star represents fluorophore (Cy5). **(F)** Representative gel images showing RAD51 filament stabilisation by full-length RAD51AP1 (wild type or mutants). Cropped images are taken from the same gel. **(G)** Schematic of oligonucleotide strand exchange assay. Star represents fluorophore (Cy5). **(H)** Representative gel images showing the stimulation of RAD51-driven strand exchange by full-length RAD51AP1 (wild type or mutants). Cropped images are taken from the same gel. **(I)** CryoEM map of the Ca^2+^-ATP bound RAD51 filament in the presence of RAD51AP1 with an AlphaFold3 model of the complex of two RAD51 (yellow and blue), C59 (magenta) and ssDNA (pink). Density outside the RAD51 filament is shown as a mesh.

To corroborate these findings in the context of full length proteins, we next tested the ability of full-length RAD51AP1 to stabilise RAD51 filaments in a nuclease protection assay (Carver et al., 2024; Taylor et al., 2015). When RAD51 forms a filament around fluorescently labelled ssDNA, the ssDNA is protected from nucleolytic digestion; more stable filaments are more resistant to nuclease digestion (**Fig. 2E**). Under our experimental conditions, RAD51 or RAD51AP1 alone offered little protection against nuclease digestion (**Fig. 2F**). By contrast, when the two proteins were added together, ssDNA protection increased substantially in a manner that was dependent on the concentration of RAD51AP1. Mutating either site reduced the effects of nuclease protection, although Site-C mutation had a larger effect, consistent with the results of our negative stain electron microscopy experiments.

We next tested the effect of RAD51AP1 on strand exchange. We used a 60 bp double-stranded DNA with homology to a 116 nt ssDNA (**Fig. 2G**). Pre-incubating RAD51 with ssDNA allows for the formation of RAD51-ssDNA filaments, which can pair with homologous dsDNA, invade the duplex, then transfer strands to form a heteroduplex molecule that has 60 bp of dsDNA and 56 nt of ssDNA. Fluorescently labelling the complementary strand on the dsDNA allowed the detection of this product following electrophoresis (**Fig. 2G**). Indeed, under these experimental conditions, strand exchange was enhanced by RAD51AP1 in a concentration-dependent manner, with only the Site-C mutant displaying a noticeable defect in this assay (**Fig. 2H**).

### CryoEM structure of the Ca^2+^-ATP RAD51-ssDNA filament bound to full length RAD51AP1

To understand how the two RAD51 binding sites of RAD51AP1 enhance RAD51 binding synergistically and how RAD51AP1 stabilises RAD51-ssDNA filaments, we sought to determine the structure of RAD51AP1 in complex with RAD51-ssDNA filaments using cryo electron microscopy.

Consistent with its binding across multiple RAD51 molecules, RAD51AP1 binding tends to induce filament clustering and bundling, preventing structural studies using single particle analysis. We therefore first attached RAD51-ssDNA filaments stabilised by Ca^2+^ to a carbon surface on EM grids, then incubated the grids with RAD51AP1 before flash freezing the grids, which resulted in well-separated filaments suitable for single particle analysis (**Fig. S2A**). Image analysis and data processing led to a 3D reconstruction of the RAD51 filament at an overall resolution of 3.1 Å (**Fig S2A**). After selecting particles in which we observed extra density on the outside of the filament, we obtained a reconstruction of the filament at an overall resolution of 3.1 Å (**Fig. 2I**, **Table 1, Fig. S2**). The reconstruction has well resolved density for ssDNA bases, ATP bound in-between RAD51 monomers and side chains of RAD51, allowing us to build a structural model of RAD51-ssDNA filament (**Fig. S2B-C**). There is additional density on the surface of some of the RAD51 monomers that could correspond to RAD51AP1 (**Fig. 2H**). Using AlphaFold3 (Abramson et al., 2024) to model a complex between a RAD51 dimer and RAD51AP1-C59 (**Fig. S2D**), we obtained a structural model that allowed the residues 329-349 of RAD51AP1 to be fitted into the extra density (**Fig. 2I**).

**Table 1.** cryoEM data collection and model statistics.

|  | RAD51 +<br>RAD51AP1 | RAD51 | RAD51 | RAD51 +<br>RAD51AP1 (C29) |
| --- | --- | --- | --- | --- |
| nucleotide | CaATP | MgATP | MgADP | MgATP |
| Microscope | Titan Krios |  |  |  |
| Voltage (keV) | 300 |  |  |  |
| Detector | K3 (Gatan) |  |  |  |
| Magnification | 81,000 |  |  |  |
| Defocus range | -3.0 to -0.9 |  |  |  |
| Frames/movie | 40 | 50 |  |  |
| Pixel size | 1.08 | 1.08 |  |  |
| Electron dose | 42 | 53 |  |  |
| Exposure (s) | 3.85 | 4.6 |  |  |
| Movies | 5001 | 3096 |  | 3233 |
| Picked particles | 2.4 million | 5.6 million |  | 6.1 million |
| Final particles | 137,283 | 242,944 | 229,610 | 97,029 |
| Processing<br>method | Single particle<br>analysis | Helical refinement | Helical refinement | Single particle<br>analysis |
| Resolution | 3.1 | 3.3 | 3.6 | 3.0 |
| *Real-space*<br>*refinement* |  |  |  |  |
| Composition: |  |  |  |  |
| Non-H atoms | 15,950 | 14,932 | 14,791 | 16,097 |
| Residues |  |  |  |  |
| Protein | 1,997 | 1,865 | 1,919 | 2015 |
| Nucleotide | 21 | 20 | 0 | 20 |
| Ligands | 21 | 19 | 18 | 17 |
| Cross-<br>correlation (CC,<br>mask) | 0.8268 | 0.8159 | 0.7813 | 0.7973 |
| Bonds (rmsd) |  |  |  |  |
| Lengths (Å) | 0.0021 | 0.0022 | 0.0044 | 0.0025 |
| Angles (°) | 0.47 | 0.42 | 0.75 | 0.47 |
| MolProbity<br>score | 1.28 | 1.24 | 1.68 | 1.29 |
| Clash score | 5.17 | 4.62 | 6.84 | 4.5 |
| Rotamer<br>outliers (%) | 0.38 | 0.54 | 0.8 | 0.5 |
| Ramachandran<br>plot |  |  |  |  |
| Outliers (%) | 0.0 | 0.05 | 0.16 | 0.05 |
| Allowed (%) | 2.02 | 1.74 | 4.14 | 2.25 |
| Favoured (%) | 97.98 | 98.21 | 95.7 | 97.7 |
| B factors |  |  |  |  |
| mean | 90.9 | 75.09 | 76.08 | 105.34 |
| stdev | 30.39 | 29.98 | 25.22 | 36.37 |

### cryoEM structures of RAD51-ssDNA filaments in the presence of Mg^2+^-ATP and Mg^2+^- ADP

Since the ATPase activity of RAD51 plays a key role in HR and RAD51’s ATPase activity is inhibited by Ca^2+^ (Stark et al., 2002), we optimised conditions including DNA sequences, RAD51 to DNA ratios, and salt concentrations that allow sufficiently stable RAD51-ssDNA filament formation in the presence of Mg^2+^ instead of Ca^2+^. We observed filaments with a range of helical parameters, highlighting the heterogeneity of these filaments. We refined two distinct 3D reconstructions to 3.6 and 3.3 Å resolution (**Table 1, Fig. S3A**). The two filaments have a rise of 18.5 Å and 16.4 Å, respectively, and a corresponding helical twist of 52.9° and 55.1°, yielding respective helical pitches of 126 Å (6.8 protomers per turn) and 107 Å (6.5 protomers per turn) (**Fig. 3A, Table S1**).

**Figure 3.**
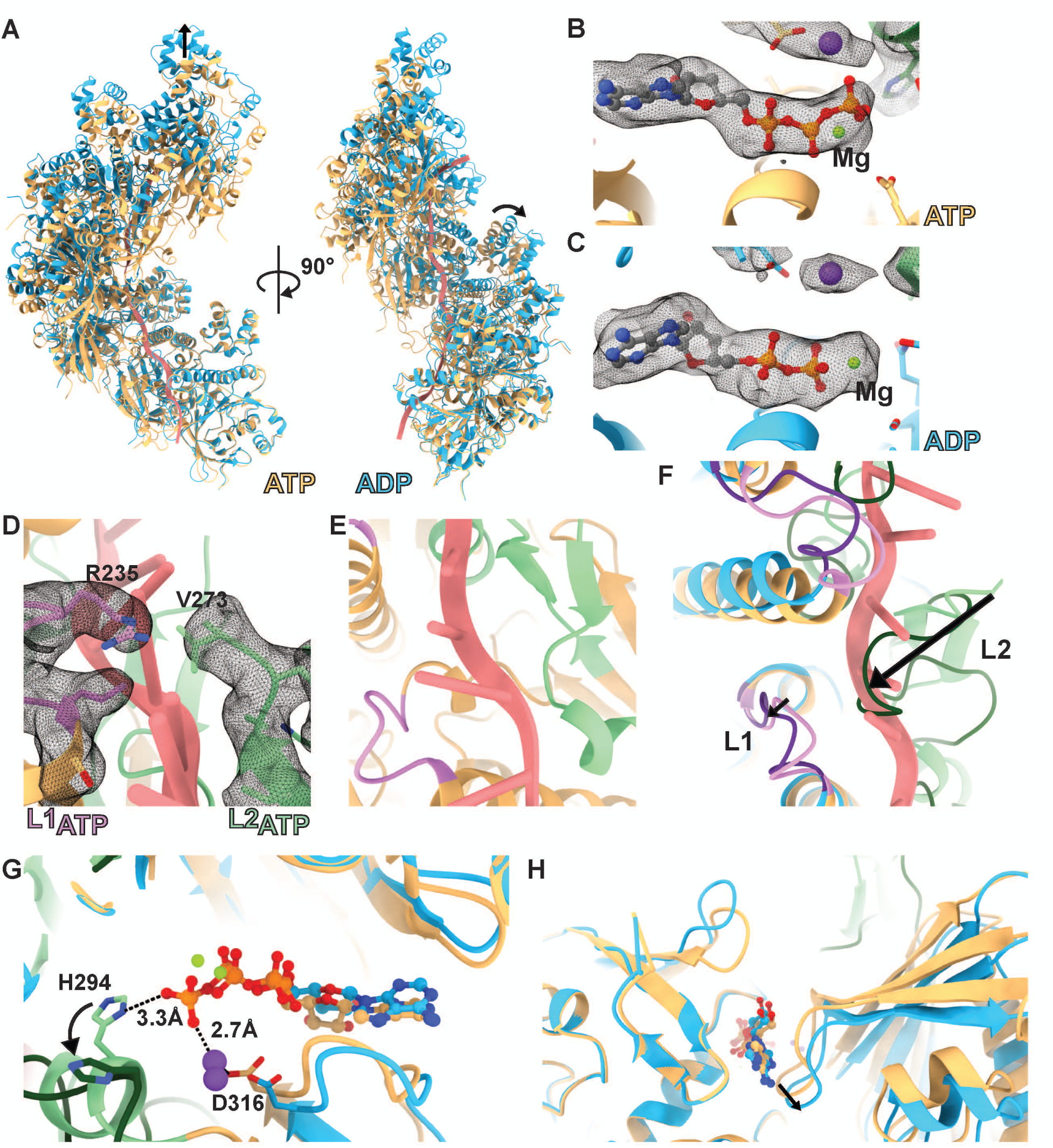
CryoEM structures of Mg^2+^-ATP and Mg^2+^-ADP bound RAD51 filaments. **(A)** Comparison of the Mg^2+^-ATP bound RAD51 filament (yellow) with the more relaxed, Mg^2+^-ADP bound filament (blue). **(B)** RAD51 nucleotide binding pocket in the ATP-bound state. **(C)** RAD51 nucleotide binding pocket in the ADP-bound state. **(D)** L1 and L2 loops of the ATP bound RAD51 filament separate the bound DNA bases into triplets. **(E)** Interaction of the β-hairpin and the α-helix of the L2 loop with the DNA backbone. **(F)** Overlay of the ATP and ADP bound filaments showing that part of the L2 loop in the ADP-bound state occupying the DNA triplet binding site in the ATP-bound state. **(G)** Overlay of the nucleotide binding sites showing that H294 coordinates the γ-phosphate of ATP and swings away from the nucleotide when ADP is bound. **(H)** Conformational changes at the RAD51 dimer interface (arrow) between ATP and ADP states.

Detailed inspections of the two reconstructions revealed that the filament with a shorter pitch has Mg^2+^-ATP bound while the one with a larger pitch has Mg^2+^-ADP bound, suggesting that in our samples, some ATP has been hydrolysed, resulting in ADP-bound filaments (**Fig. 3B-C**). The ATP-bound structure is similar to those obtained using Mg^2+^-AMP-PNP or Ca^2+^- ATP (Appleby et al., 2023b; Xu et al., 2017). The ssDNA is organised into stacked triplets intercalated by the conserved L1 and L2 loops of RAD51, specifically R235 of L1 and V273 of L2 (**Fig. 3D**). Further, the β-hairpin and the α-helix within the L2 region are also involved in DNA binding (**Fig. 3E**). In the ADP-bound filament, we do not observe clear density for ssDNA, consistent with ADP-bound filaments having weaker DNA binding, DNA has dissociated, either fully or partially (van Mameren et al., 2009).

Comparing the two structures, we observe the unfolding of the L2 region, losing its α-helix adjacent to the γ-phosphate, as well as the β-hairpin (**Fig. 3E-F**). In the ATP-bound structure, the γ-phosphate helps to coordinate the H294 side chain at the end of the small α-helix of L2 (**Fig. 3G**). Loss of the γ-phosphate leads to the rotation of H294 side chain and subsequent reorganisation of the nucleotide-binding pocket. Consequently, the α-helix becomes unfolded and the L2 loop is re-configured with the β-hairpin becoming a random coil (**Fig. 3E-F**). The L1 is also positioned slightly away from the ssDNA. These changes in L1 and L2 result in a reduction in DNA binding. In our structure, the L2 tip now occupies the ssDNA position (**Fig. 3F**). However, it is unclear if this repositioning contributes to DNA dissociation or is due to availability of the space vacated by the release of ssDNA. We also observe changes at the the protomer interace (**Fig. 3H**), which becomes looser in the ADP-bound form, resulting in the smaller twist angle and higher helical rise between two protomers (**Fig. 3H, 3A**).

### cryoEM structures of the Mg^2+^-ATP bound RAD51-ssDNA in complex with the RAD51AP1 C-terminal domain

The quality of density corresponding to RAD51AP1 is limited, probably due to the heterogeneity arising from the two RAD51 binding sites across two RAD51 monomers, and Site-N having a weaker affinity for RAD51. Given that mutation of Site-N had limited effect on binding (**Fig. 1B**, **Fig. 2D**), we designed a simpler construct that lacks Site-N but contains the major binding site Site-C (C29, C-terminal 29 residues). We reasoned that C29 would bind more uniformly to every RAD51 protomer in the filament, improving occupancy and homogeneity of the samples. The structure was resolved to 3.0 Å resolution (**Fig. S4A**), clearly revealing Mg^2+^-ATP, ssDNA (**Fig. S4B-C**), and density corresponding to RAD51AP1 (**Fig. 4A, B**).

**Figure 4.**
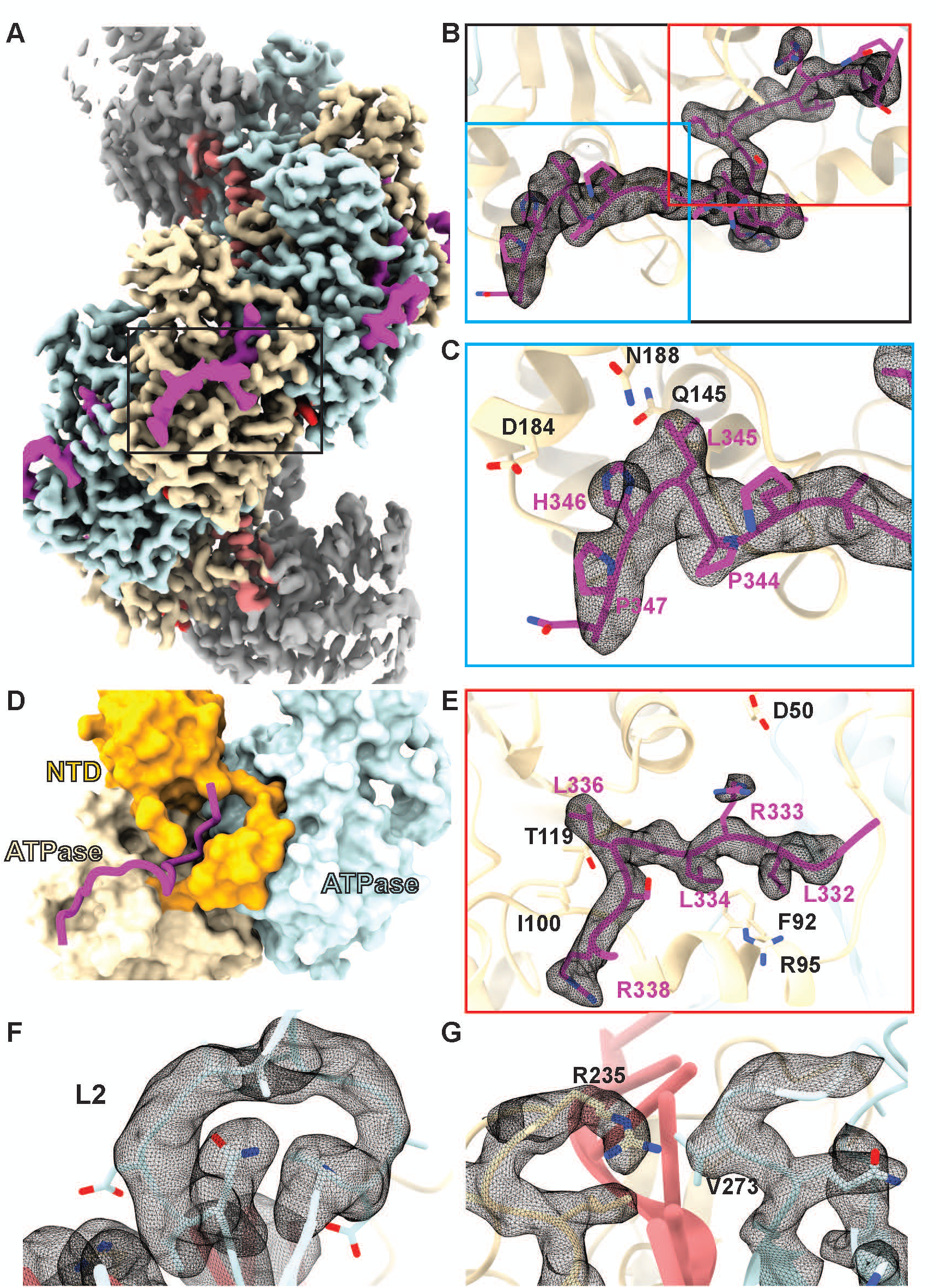
CryoEM structure of the Mg^2+^-ATP bound RAD51 filament in complex with the RAD51AP1 C29 fragment. **(A)** CryoEM map of the RAD51 (yellow and blue) filament on ssDNA (pink) bound by C29 (magenta). **(B)** Closeup view of the density corresponding to RAD51AP1. **(C)** Closeup view of the density corresponding to RAD51AP1 Site-C. **(D)** Surface representation of RAD51 showing RAD51AP1 binding in a groove created between the N-terminal domain and the ATPase domain of RAD51. **(E)** Closeup view of the density corresponding to RAD51AP1 Site-M. **(F)** Fully resolved density of the L2 loop of RAD51 when bound to RAD51AP1 C29. **(G)** Residues of the L1 and L2 loop that separate the bound DNA bases into triplets.

The filaments have a helical twist of 55.8° and a rise of 15.9 Å, giving rise to a helical pitch of 103 Å and 6.5 protomers per turn (**Table S1**). This is similar to the Ca^2+^-ATP filament bound to full-length RAD51AP1 (104 Å and 6.4 protomers per turn), further supporting the notion that C29 contains the major sites responsible for RAD51 filament modulation. The previously identified H346 in Site-C inserts into a shallow groove on the RAD51 surface, interacting with the side chain of D184 and the main chain of Q145 of RAD51 (**Fig. 4C**), explaining its crucial roles in binding (**Fig. 1B-D**). The adjacent L345 also inserts itself into the same shallow groove. Interestingly, both in the structure of full-length RAD51AP1 in complex with Ca^2+^-ATP-RAD51-ssDNA (**Fig. 2I**) and that of the C29-bound Mg^2+^-ATP-RAD51-ssDNA, we observe continuous density located N-terminal to Site-C (**Fig. 4B**). Indeed, the density resides in a groove created between the RAD51 N-terminal helical bundle and the N-terminal helix (**Fig. 4D**). This site contains the ^332^LRLGLSR^338^ motif, hereafter referred to as Site-M, which inserts itself into a pocket lined by largely hydrophobic residues including RAD51-F92, R95 (surrounding RAD51AP1 L334), I100, and T119 (surrounding RAD51AP1 L336) (**Fig. 4D-E**). Interestingly, previous yeast two-hybrid analysis demonstrated that L336Q (L319Q in (Kovalenko et al., 2006; Wiese et al., 2007)) severely disrupts the interaction between RAD51 and RAD51AP1, and this was confirmed in binding assays with purified proteins (Wiese et al., 2007), consistent with our structure here. Charge interactions between RAD51AP1-R333 and RAD51-D50 accommodate R333 in the pocket and help to stabilise the RAD51 N-terminus, now with density visible for residues 19-20 above Site-M (**Fig. 4D**); these residues were absent in the structures without RAD51AP1 bound, suggesting that RAD51AP1 binding can also stabilise the RAD51 N-terminus.

Looking closer into the structural changes within RAD51, we observe that upon RAD51AP1 binding, the binding pocket, especially around Site-M, remodels to accommodate the peptide. Interestingly, we now observe density for the entire L2 region (**Fig. 4F**), for which the density of the tip was missing in the RAD51-ssDNA structures, suggesting L2 adopts a more rigid conformation upon RAD51AP1 binding. RAD51AP1 binding induces changes such that the β-hairpin at the base of the L2 loop is brought closer to DNA, strengthening its interactions with DNA, and subsequently stabilising the tip of the L2, re-enforcing the interactions between V273, the DNA base and R235 of L1 (**Fig. 4G**).

### RAD51AP1 modulates RAD51 DNA binding, filament stabilisation, and strand invasion via three distinct sites

To corroborate the structural observations and the functional relevance of the interactions, especially the role of Site-M, we investigated the effects of RAD51AP1 binding on RAD51 HR functions using mutagenesis and biochemical methods. Given that nucleotides bind in-between two RAD51 protomers, we tested whether the nucleotide influences RAD51AP1 binding to RAD51. The omission of ATP and Mg^2+^ in the pull-down buffer led to a complete loss of RAD51 binding (**Fig. 5A-B**), supporting the model that RAD51AP1 binds across two RAD51 protomers. This is further supported by the lack of binding to the RAD51-F86E mutant, which is severely defective for oligomerisation (**Fig. S5A**)(Esashi et al., 2007).

**Figure 5.**
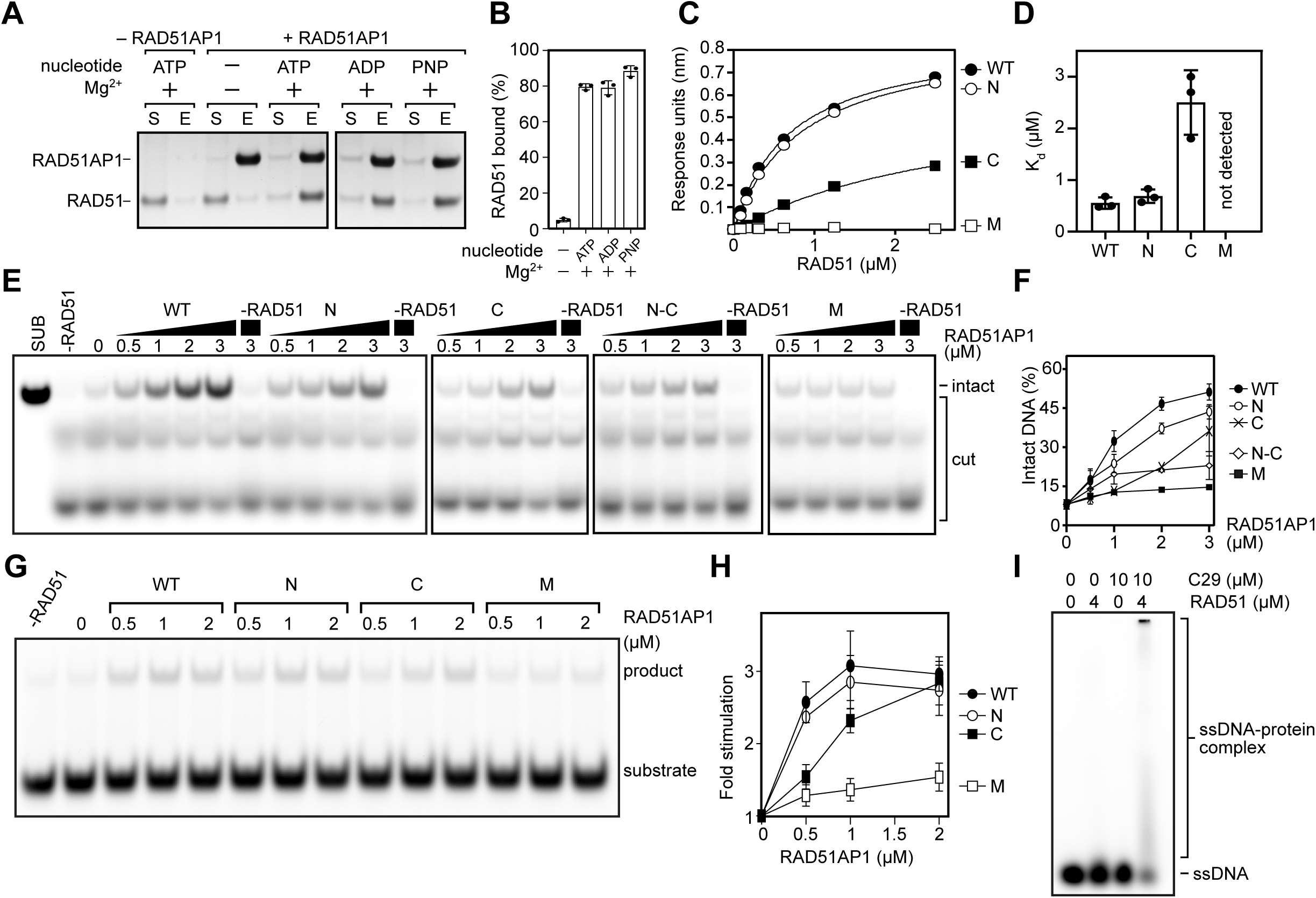
Integrity of Site-M is required for RAD51AP1 activities. **(A)** Pull-down experiments of full-length RAD51AP1 (6xHis-tagged) and RAD51 in the presence of the indicated co-factors. **(B)** Quantification of **(A)**. Averages are shown with error bars depicting the standard deviation (n = 3). PNP, AMP-PNP. **(C)** Binding curves of RAD51AP1-C59 (wild type or mutants) obtained from bio-layer interferometry. A representative result is shown. **(D)** Kd values for the binding of RAD51AP1-C59 and RAD51, determined by bio-layer interferometry. Averages are shown with error bars depicting standard deviation (n = 3). **(E)** Representative gel images showing RAD51 filament stabilisation by full-length RAD51AP1 (wild type or mutants). **(F)** Quantification of **(E)**. Averages are shown with error bars depicting standard deviation (n = 3). **(G)** Representative gel image showing the stimulation of RAD51-driven strand exchange by full-length RAD51AP1 (wild type or mutants). **(H)** Quantification of **(G)**. Averages are shown with error bars depicting standard deviation (n = 3). **(I)** Representative gel image showing the electrophoretic mobility of RAD51-bound 10 nt ssDNA, with and without RAD51AP1-C29.

**Figure 6.**
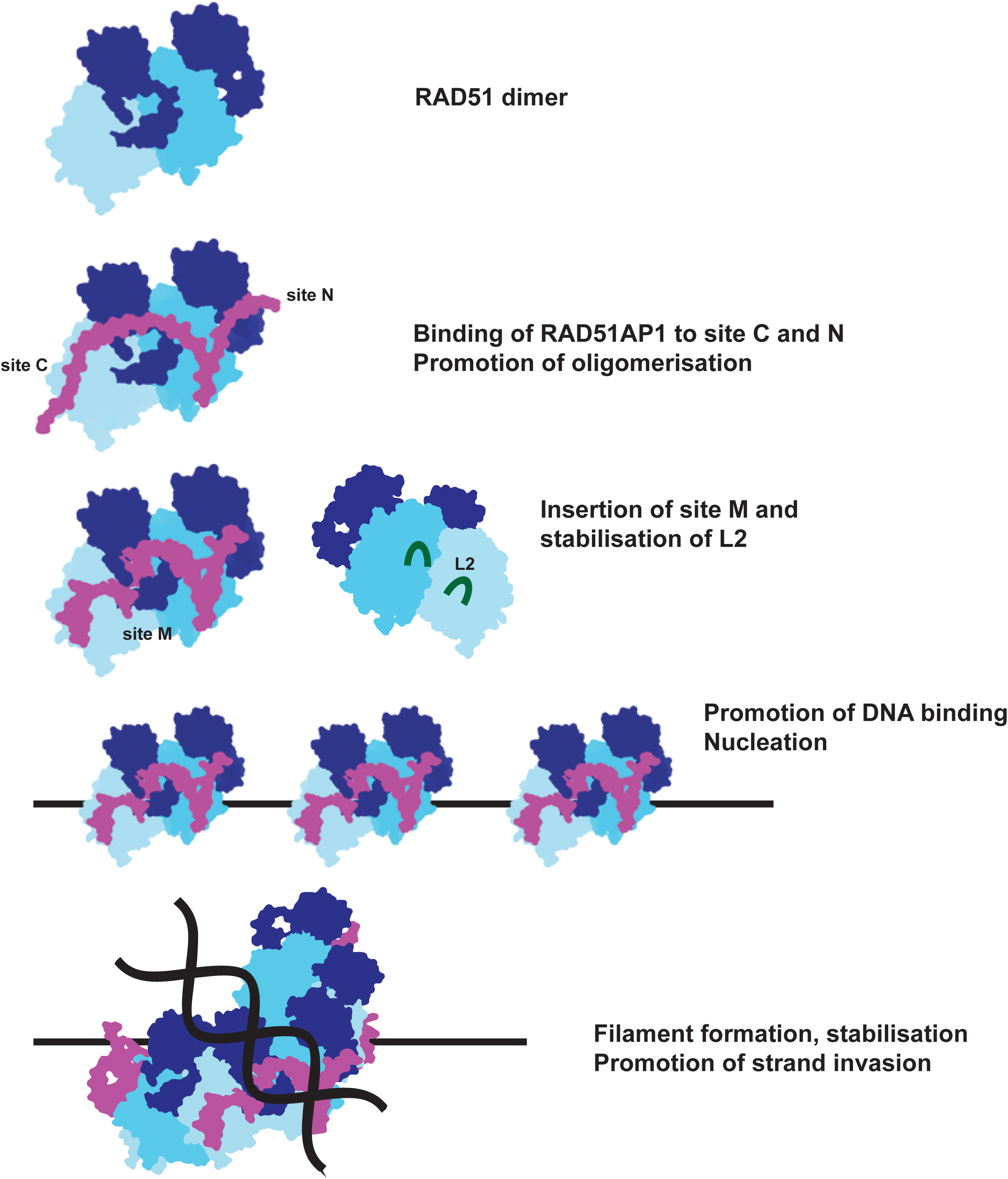
A proposed model for RAD51AP1 in modulating RAD51 activities. Cartoon model of RAD51AP1 binding to the RAD51 dimer, using Site-N and Site-C, promoting oligomerisation. Site-M then inserts and strengthens the interface and promotes the ordering of the L2 loop, stabilising DNA binding and filament nucleation. Once the filament is assembled, the RAD51AP1 bound filament will be more stable and efficient in strand exchange.

Site-M binds at the dimer interface and its binding induces conformational changes and remodels the filament, suggesting a crucial role in both binding and remodelling. We mutated key residues in Site-M (RLGL to AAGA) and characterised its effects on RAD51 binding, nuclease protection and strand invasion. We first examined the effects of Site-M mutation in the context of the simplified C59 construct, which can bind RAD51 almost as well as full-length RAD51AP1 (**Fig. 2D**). Through BLI experiments, we found the Site-M mutant to be severely defective in RAD51 binding, even more so than the Site-C mutant (**Fig. 5C-D**). Consistent with the associated binding defect, mutation of Site-M rendered nuclease protection by RAD51AP1 ineffectual (**Fig. 5E-F**). The C59 construct was also able to achieve maximal RAD51 filament stabilisation in the nuclease protection assay, albeit at higher concentrations compared to full-length RAD51AP1 (**Fig. S5D**). Similarly to the full length protein, the Site-M mutant in the C59 context was defective for nuclease protection (**Fig. S5D**). Moreover, the full-length Site-M mutant was severely impaired in the stimulation of RAD51-driven DNA strand exchange **(Fig. 5G-H**). These results are in agreement with our structural observations indicating that the C-terminal region of RAD51AP1, and in particularly Site-M, constitutes the critical module responsible for RAD51 regulation.

Although mutation of Site-N had no obvious effect on RAD51 binding (**Fig. 1B-D** and **Fig. 2D**), it nevertheless resulted in a reduction in filament numbers, suggesting a defect in filament nucleation; this is consistent with its importance in tethering across two protomers. By contrast, mutation of Site-C reduced binding, and as expected for a physical binding defect, the impairment in filament stabilisation could be overcome by increasing the concentration of RAD51AP1 (**Fig. 5E-H**).

To investigate if RAD51AP1 could indeed promote RAD51 association with DNA as suggested by the structure, especially due to the changes in the L2 region (**Fig. 4F-G**), we measured RAD51 binding to a short ssDNA (10 nt) (**Fig 5I**). Since each RAD51 can bind to three nucleotides, 10 nt could accommodate a maximum of 3 RAD51 molecules; we reasoned that this would prevent stable filament formation and therefore allow for the measurement of DNA binding *per se,* rather than binding and subsequent filament formation. As expected, the binding is weak with significant unbound DNA remaining. However, the amount of shifted DNA increased in the presence of C29, suggesting that C29 enhances the initial association of RAD51 with ssDNA.

## Discussion

### ATP hydrolysis relaxes rather than compresses RAD51 filaments

Thus far, most high resolution RAD51 filament structures were determined without hydrolysable ATP, either Ca^2+^-ATP or Mg^2+^-AMPPNP, and all showed similar structures with a helical pitch around 105 Å and 6.4 protomers per turn (**Table1. S1**). Our Mg^2+^-ATP bound RAD51-ssDNA filament displayed a slightly more relaxed filament (107 Å and 6.5 protomers per turn). The ATP-bound filament has been shown to form on pre-synaptic (ss) and post-synaptic (ds) DNA in a similar fashion, with DNA stretched between B-form nucleotide triplets, the ATP bound RAD51-ssDNA is therefore ready for pairing with complementary strand. Single molecule experiments showed that dsDNA is stretched upon addition of RAD51 and ATP, consistent with RAD51-ATP binding stretching the DNA (Hilario et al., 2009). Recently published structures of the Ca^2+^-ADP bound (Appleby et al., 2023a) and Mg^2+^-ADP bound filaments (Luo et al., 2023) showed a more extended filament conformation, although the Mg^2+^-ADP structure had two filaments intertwined (Luo et al., 2023). In that structure, each single filament is extended to a helical pitch of ∼145 Å (Appleby et al., 2023a; Luo et al., 2023). Our Mg^2+^-ADP bound structure has a helical pitch of 126 Å, almost identical to that of Ca^2+^- ADP filaments. These structural data therefore reveal that ATP hydrolysis relaxes RAD51 filaments, contrary to previous single molecule experiments that suggested that ATP hydrolysis contracts the filaments (Robertson et al., 2009). However, that single molecule study was based on DNA length measurements. Indeed, another single molecule study monitoring both RAD51 binding and DNA length showed that ADP-bound RAD51 can bind to dsDNA but does not extend it (Hilario et al., 2009); in comparison to ATP-bound RAD51 filaments where DNA is stretched, the DNA appeared to be more compact in the ADP-bound filaments. Intriguingly, DNA stretching is shown to slow down RAD51 dissociation post-hydrolysis (van Mameren et al., 2009). These seemingly conflicting results can be reconciled by the conformational changes observed in our ADP-bound filament structure. When the DNA is further stretched, the extended ADP filament could enable the L1/L2 loops to intercalate between stretched bases, therefore reducing the dissociation from DNA post-hydrolysis as observed (van Mameren et al., 2009). RAD51-ATP binding stretches dsDNA, enabling stable interactions between L1/L2 loops and DNA. ATP hydrolysis now further stretches the RAD51 filaments, breaking the interactions with DNA, enabling the stretched DNA to return to the preferred B-form, resulting in DNA compaction observed in ^25^. Interestingly, the Ca^2+^-ATP bound ssDNA filaments, either alone or in complex with RAD51AP1 or TR2 (the C-terminal domain of BRCA2), have an even smaller helical pitch (103-104 Å) and higher twist (6.4 protomers per turn), suggesting an even more tightly packed filament compared to Mg^2+^-ATP (107 Å and 6.5 protomers per turn). These data suggest that many RAD51 modulators stabilise the RAD51 filaments by tightening the filaments, whereas ATP hydrolysis relaxes the filaments, allowing DNA to resume the stable B-form. This could be important for processes post-synapsis, especially those involving modulators such as RAD54 and RTEL but also RAD51AP1 (Modesti et al., 2007), which bind to B-form dsDNA (Ristic et al., 2001).

### Structural basis of RAD51-DNA association in the presence of ATP

RAD51-filaments are stabilised by DNA interactions via the L1 and L2 loops, the protomer-protomer interactions mediated by the ATPase domains, and the N-terminal domain, which bridges the two ATPase domains (**Fig. 4D**). In the Mg^2+^-ATP and Mg^2+^-ADP bound RAD51 filament structures presented here, we observe major conformational changes in the L2 regions (**Fig. 3**). Recent structural studies of RAD51-ssDNA in the presence of Ca^2+^-ATP or Ca^2+^-ADP revealed that the γ-phosphate coordinates a second metal ion which is absent in the ADP-bound form, and the authors proposed that this second ion contributed to the folding of L2 (Appleby et al., 2023a). Interestingly, we observed density corresponding to the second ion in both structures (**Fig. 3B-C**). The second ion interacts with both γ-phosphate and D316, which does not change in the ADP-bound structure. The density corresponding to the second ion in the ADP structure is weaker, which could result from reduced affinity due to the loss of γ-phosphate, or because the resolution of the reconstruction is lower. However, H294 in the short α-helix of L2 does not interact with the second metal ion in the ATP-bound state, and its re-orientation seems to be entirely due to the loss of γ-phosphate, suggesting that the second ion might not affect the folding state of this α-helix (**Fig. 3G**). An earlier crystal structure of RadA also observed a second ion at this position, where it was proposed to play a role in coordinating and polarising the catalytic water (Wu et al., 2005). It is possible that the second ion plays a similar catalytic (as opposed to structural) roles in RAD51. Indeed, K+ or Na+ both can stimulate ATPase activity (**Fig. S6A**).

### RAD51AP1 binds to RAD51 using a unique binding mode

Our data here reveal that RAD51AP1 binds across two RAD51 protomers in the RAD51-ssDNA filament via 3 distinct sites. Site-M is the most important site as the mutant does not bind to RAD51 and is unable to promote filament stabilisation or stimulate strand exchange. We propose that Site-N mainly plays a role in filament nucleation via its crucial role in facilitating the binding across two protomers. However, we postulate that Site-N is less important for other aspects of filament stabilisation, and this is supported by our biochemical assays — where its defects were milder than the Site-C/M mutants — and the structural similarities of the C29-bound RAD51 filament — which completely lacks Site-N — and the full-length RAD51AP1-bound filament. Since Site-M inserts itself into a deep pocket on the RAD51 filament surface that is not readily accessible, we propose that Site-C, which binds at the RAD51 surface, acts as a recruitment anchor to bring RAD51AP1 onto the RAD51 filaments, which then allows Site-M to be in close proximity to the deep pocket. The subsequent insertion into the RAD51 N-terminal domain then leads to remodelling of the protomer-protomer interface, and consequently the filament itself (**Fig. 4**). Mutating Site-C severely impairs binding, but its defects are rescued at sufficiently high RAD51AP1 concentrations, consistent with its recruitment function. By contrast, mutating Site-M completely abolished all functions, supporting the notion that it is the major binding and remodelling site.

Site-M inserts into a pocket formed by the RAD51 N-terminal domain, thus stabilising the N-terminal domain and protomer-protomer interface. Furthermore, ATP/ADP in the filament is located in-between the RAD51 protomers, with the phosphates buried deep inside the filament. There is a narrow channel where the γ-phosphate could be released upon hydrolysis (**Fig. S6C**). However, this channel is now blocked by Site-M and tightened by the effect of RAD51AP1 binding on the helical parameters of the filament, suggesting that RAD51AP1 binding might also hinder γ-phosphate release, which would inhibit ATP hydrolysis and further stabilise the filament. Indeed, in the samples of RAD51-ssDNA-Mg^2+^-ATP in complex with RAD51AP1-C29, we do not observe filaments with larger helical pitches that correspond to ADP-bound filaments, consistent with inhibition of γ-phosphate release. Interestingly, previous biochemical analysis with mouse proteins has demonstrated that RAD51AP1 can reduce ATP hydrolysis by RAD51 (Su et al., 2014).

Notably, RAD51AP1 binding induces the stabilisation of the L2 loop. In RecA, I197 in L2 intercalates into the nucleotide triplets in the invading ssDNA while F203 inserts into donor dsDNA, with the aromatic side chain of F203 facilitating stacking interactions with DNA bases and maintaining the D-loop structure, as shown in the RecA strand-exchange intermediate structures(Yang et al., 2020). This region is highly conserved in RAD51, with L2 loops adopting similar conformations in our C29-bound RAD51-ssDNA filaments (**Fig. S6D**). RAD51 V293 can be superimposed with I197 of RecA while F297 in RAD51 is equivalent to RecA’s F203, suggesting similar roles. In the C29-bound structure, the stabilised L2 loop would thus also stimulate strand invasion and D-loop formation. RAD51AP1 therefore stimulates strand exchange by both stabilising filaments and optimising L2 conformation for strand invasion and D-loop stabilisation.

Comparison with other RAD51 interacting partners reveals the differences and similarities in their binding modes. Site-N is predicted to bind to RAD51 via a WVPP motif, which is reminiscent of the FQPP motif employed by C-terminal region of BRCA2 (TR2) to bind to RAD51 (**Fig S6E**) (Appleby et al., 2023b). Additional residues adjacent to FQPP in TR2 stabilises the two RAD51 monomers. RAD51AP1 uses different binding sites across two protomers (**Fig. S6E**). RAD51AP1 Site-C also shares a binding site with BRCA2 TR2, suggesting that both RAD51AP1 Site-N and Site-C would overlap with TR2 binding (**Fig. S6E**). Recent studies reveal that FIGNL1 binds at three distinct RAD51 sites. Two sites within the FRBD domain are predicted to overlap with BRCA2 BRC4 and TR2 binding sites, respectively. Uniquely, it encloses the RAD51 N-terminus in the hexamer pore formed by its C-terminal ATPase domains (Carver et al., 2024). Site-M does not overlap with the binding of TR2 or BRC4. It will be interesting to investigate if these modulators compete for binding to RAD51 (**Fig. S7**) and at what stoichiometry multiple modulators decorate RAD51 filaments during HR.

### RAD51AP1 is a versatile RAD51 modulator and remodels RAD51 filaments

Our comprehensive data here reveal the molecular mechanism of RAD51AP1 in promoting RAD51 activities in homologous recombination. We showed that via three binding sites across two RAD51 monomers, RAD51AP1 modulates almost all RAD51 activities in HR: enhancing DNA binding, filament nucleation, filament stabilisation, and strand exchange. In addition to modulating RAD51 filaments, RAD51AP1 has been proposed to act as a bridge in tethering RAD51-ssDNA filaments formed at the DSB site to the sister chromatid. Its ability to bind to TERRA and promote R-loop formation has highlighted its roles in ALT and telomere maintenance. Whether these activities are independent or act synergistically with its RAD51 modulator function, which activities are important for the different processes RAD51AP1 is involved in, and how its versatile activities are regulated, remain to be investigated.

## Resource Availability

## Acknowledgement

Initial screening of electron microscopy grids was carried out at Imperial College London Centre for Structural Biology. We acknowledge Diamond Light Source for access and support of the cryoEM facilities at the UK national electron bioimaging centre, proposal B136390, funded by the Wellcome Trust and the United Kingdom Research and Innovation Medical Research Council, and London Consortium for high resolution cryoEM (LonCEM), funded by the Wellcome Trust. We thank members of the Zhang Lab for their helpful insights and discussions. This work is funded by the Wellcome Trust to X.Z. (210658/Z/18/Z, 227769/Z/23/Z).

## Author contributions

X.Z. conceived the project, while L.K., B.A. and X.Z. designed the experiments. CryoEM sample preparation, image processing and structure determination was performed by L.K. Protein construct cloning, mutagenesis, and purification was performed by L.K. with contributions from J.Z. Biochemical assays were performed by B.A., L.K. and P.L. BLI was performed by B.A. and L.M. Bioinformatic and structural analysis was performed by L.K. and X.Z. L.K.,B.A. and X.Z. wrote the initial manuscript. Edits to the manuscript were carried out by all authors.

## Declaration of interests

Authors declare that they have no competing interest.

## Declaration of generative AI and AI-assisted technologies

AlphaFold3 server (Abramson et al., 2024) was used to generate initial structural model of RAD51AP1 which was modified and refined according to experimental data. EMReady (He et al., 2023) was used for density modification of experimental EM maps after data processing.

## Supplemental Information

**Figure S1. Related to Figure 1.**
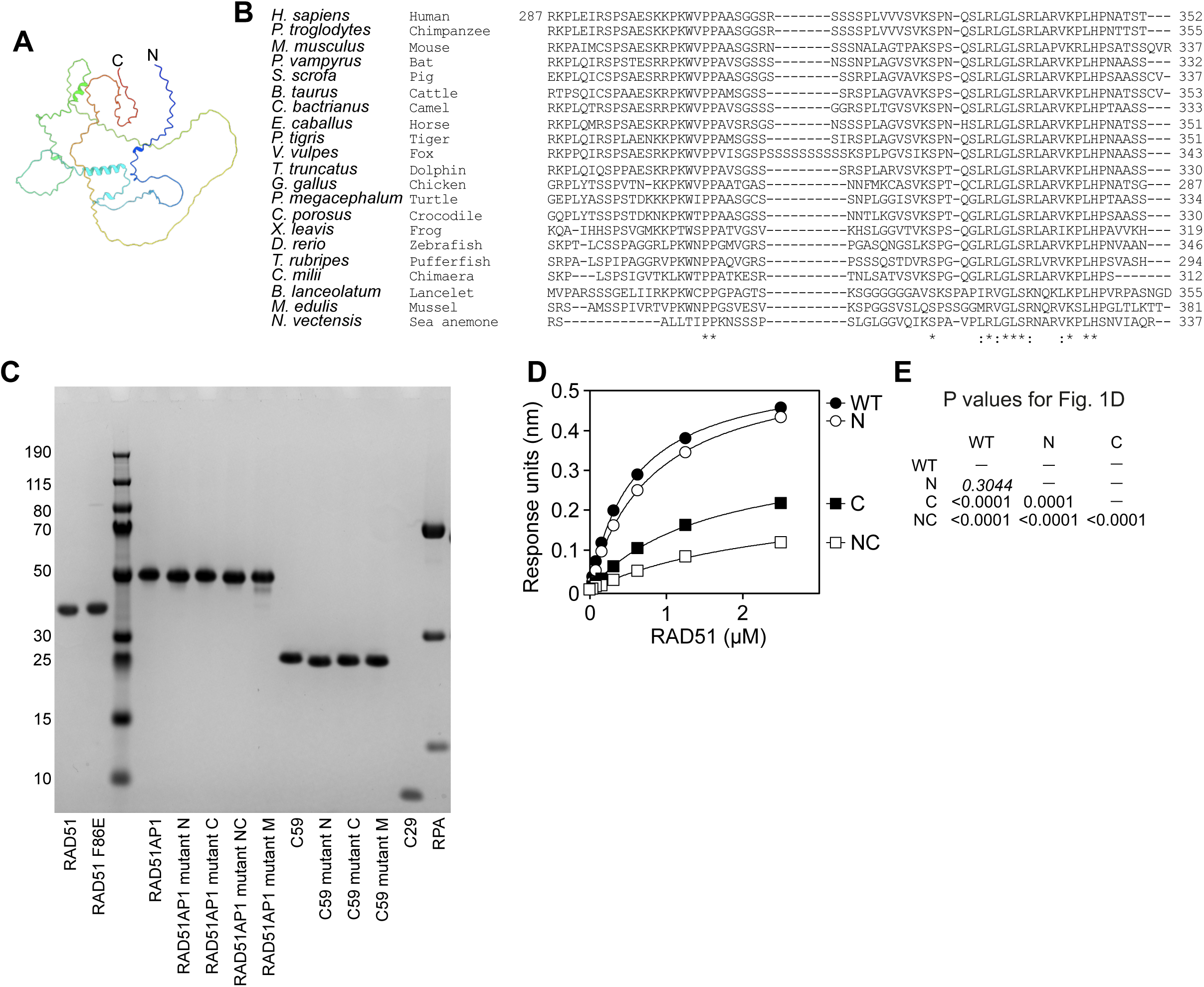
**(A)** Structural prediction by AlphaFold3 showing that RAD51AP1 is an intrinsically disordered protein. **(B)** Sequence alignment of RAD51AP1 orthologues from indicated species. **(C)** SDS-PAGE analysis demonstrating the purity of proteins used in this study. **(D)** Binding curves of full-length RAD51AP1 (wild type or mutants) obtained from bio-layer interferometry. A representative result is shown. **(E)** P value for results in Figure 1D. Statistical test was one-way ANOVA followed by Tukey’s correction for multiple comparisons.

**Figure S2. Related to Figure 2.**
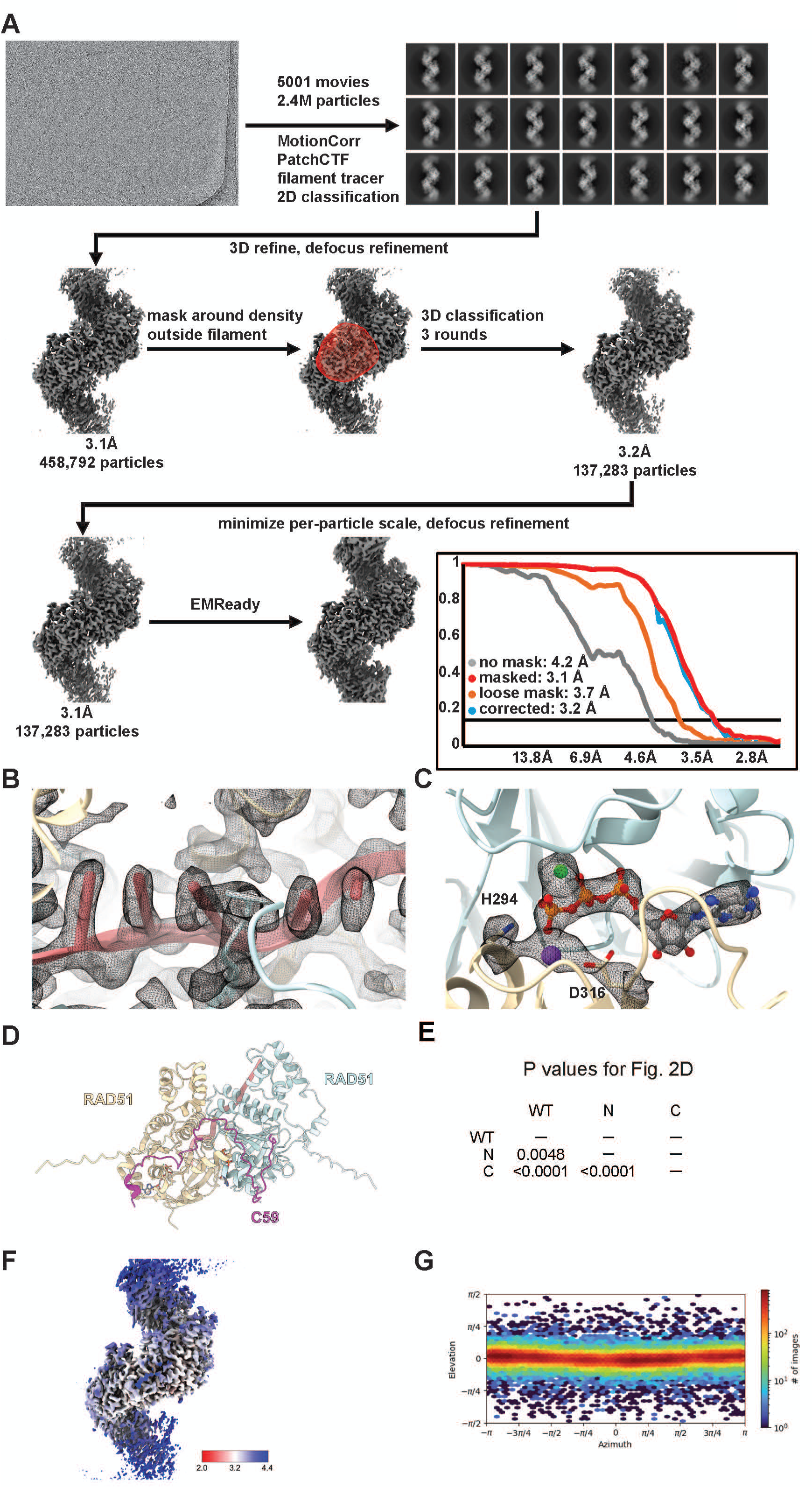
**(A)** Data processing flowchart of the Ca^2+^-ATP RAD51 filament bound to full length RAD51AP1. **(B)** Density corresponding to the DNA. **(C)** Density corresponding to ATP. **(D)** AlphaFold3 model of RAD51, ssDNA and RAD51AP1-C59. **(E)** P value for results in Figure 2D. Statistical test was one-way ANOVA followed by Tukey’s correction for multiple comparisons. **(F)** CryoEM map of the filament from Figure 2I coloured by local resolution. **(G)** Orientation distribution of the Ca^2+^-ATP filament bound by RAD51AP1.

**Figure S3. Related to Figure 3.**
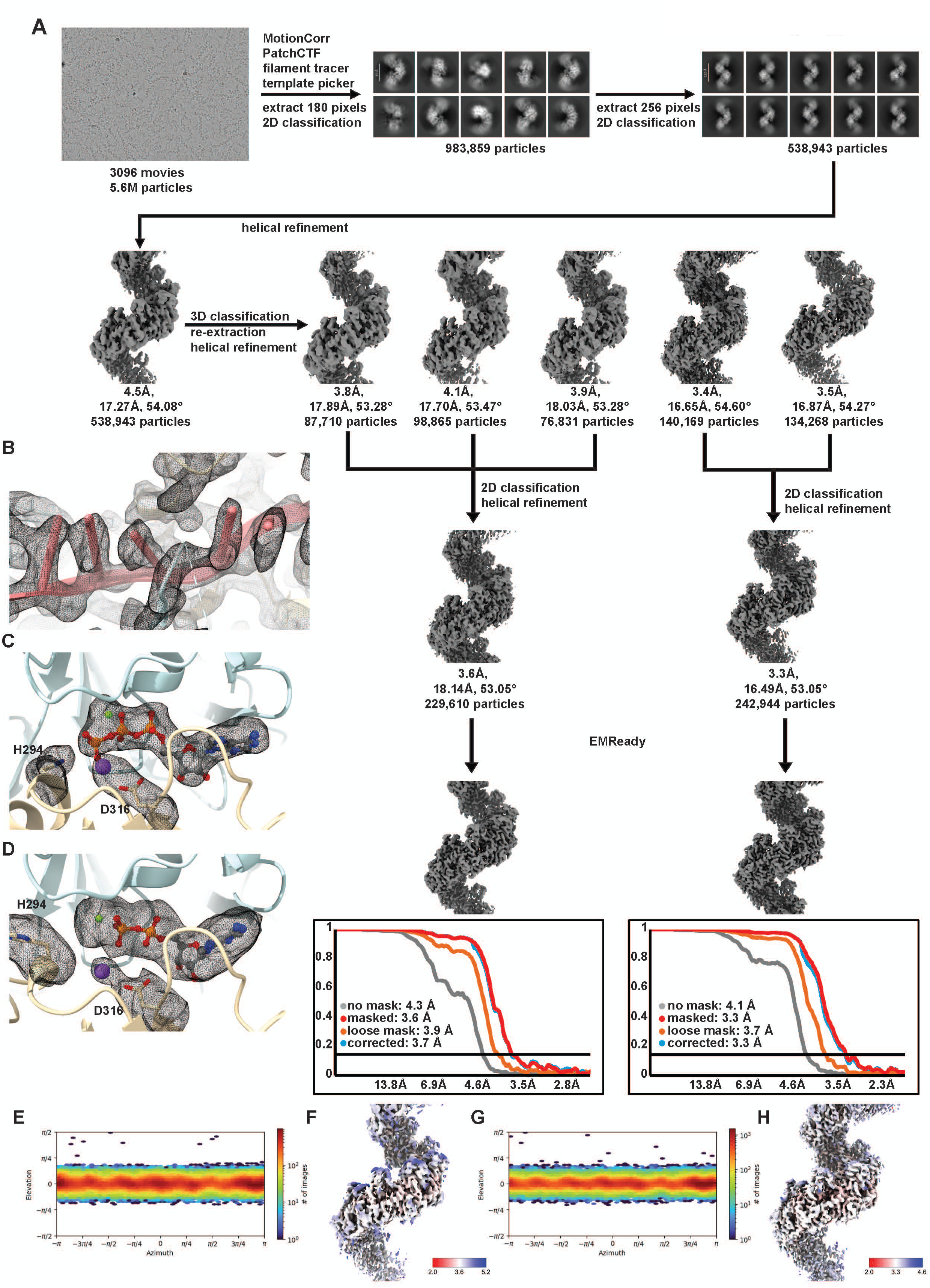
**(A)** Data processing flowchart of the RAD51-ssDNA filaments in the presence of Mg^2+^-ATP and Mg^2+^-ADP. **(B)** Density corresponding to the DNA in the ATP filament. **(C)** Density corresponding to ATP in the ATP filament. **(D)** Density corresponding to ADP in the ADP filament. **(E)** Orientation distribution of the ADP filament. **(F)** CryoEM map of the ADP filament coloured by local resolution. **(G)** Orientation distribution of the ATP filament. **(H)** CryoEM map of the ATP filament coloured by local resolution.

**Figure S4. Related to Figure 4.**
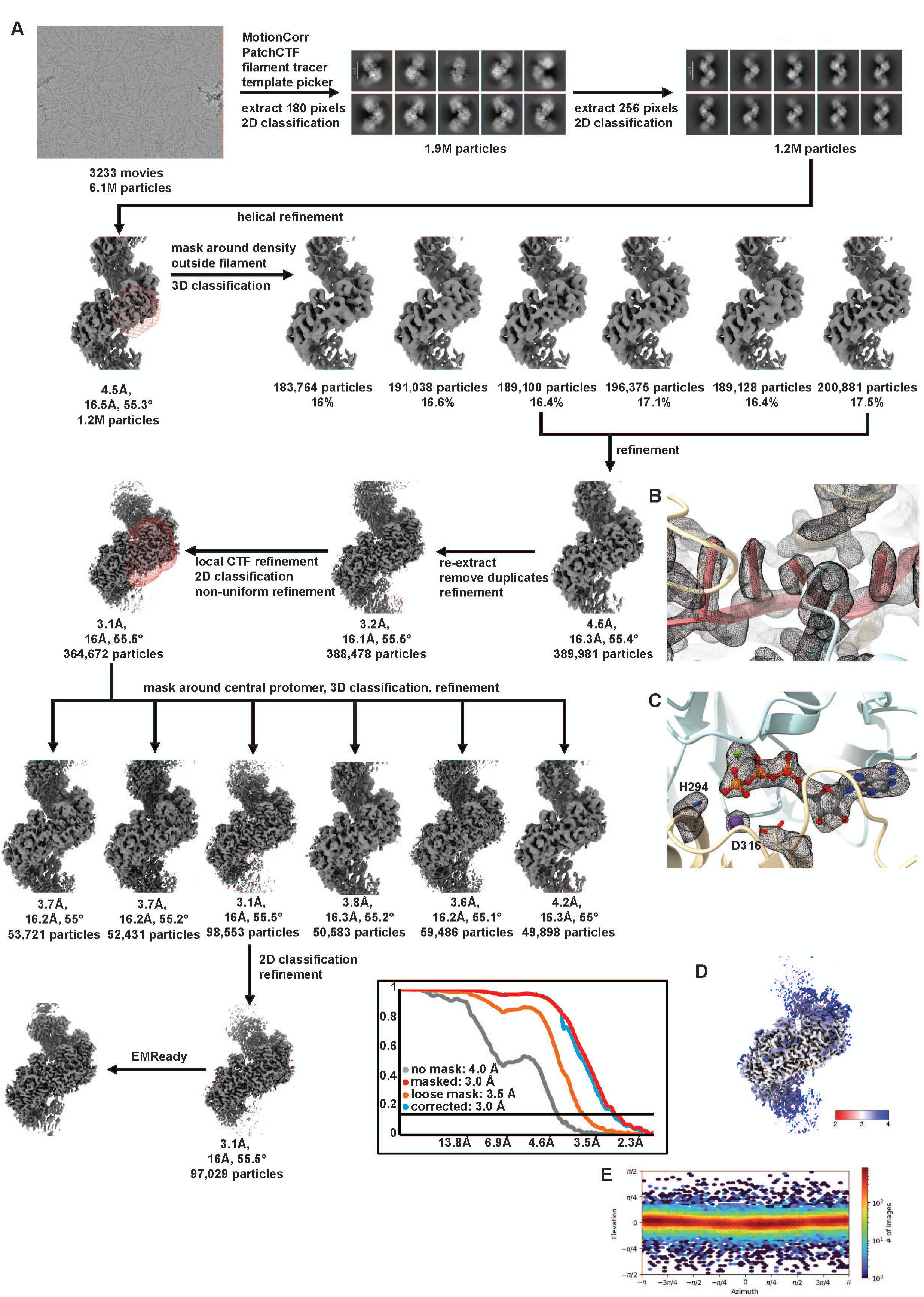
**(A)** Data processing flowchart of the Mg^2+^-ATP RAD51 filament bound by RAD51AP1 C29. **(B)** Density corresponding to the DNA in the filament. **(C)** Density corresponding to ATP. **(D)** CryoEM map of the filament coloured by local resolution. **(E)** Orientation distribution of the filament.

**Figure S5. Related to Figure 5.**
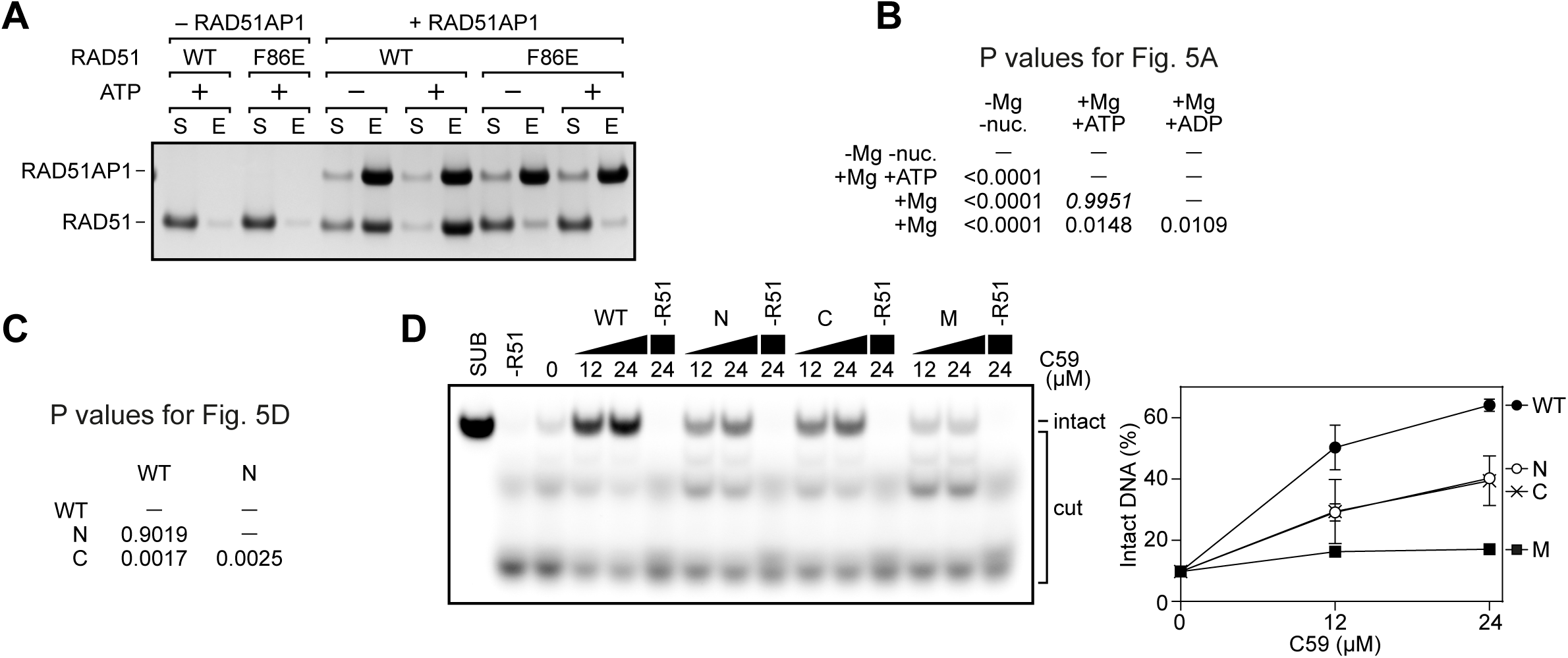
**(A)** Pull-down experiment of full-length RAD51AP1 with either RAD51 or the oligomerisation defective mutant RAD51-F86E, with and without ATP. **(B)** P values for results in Figure 5A. Statistical test was one-way ANOVA followed by Tukey’s correction for multiple comparisons. **(C)** P values for results in Figure 5D. Statistical test was one-way ANOVA followed by Tukey’s correction for multiple comparisons. **(D)** Representative gel images showing RAD51 filament stabilisation by RAD51AP1-C59 (wild type or mutants). In the quantification, averages are shown with error bars depicting standard deviation (n = 3).

**Figure S6. Related to Figure 5 and Figure 6.**
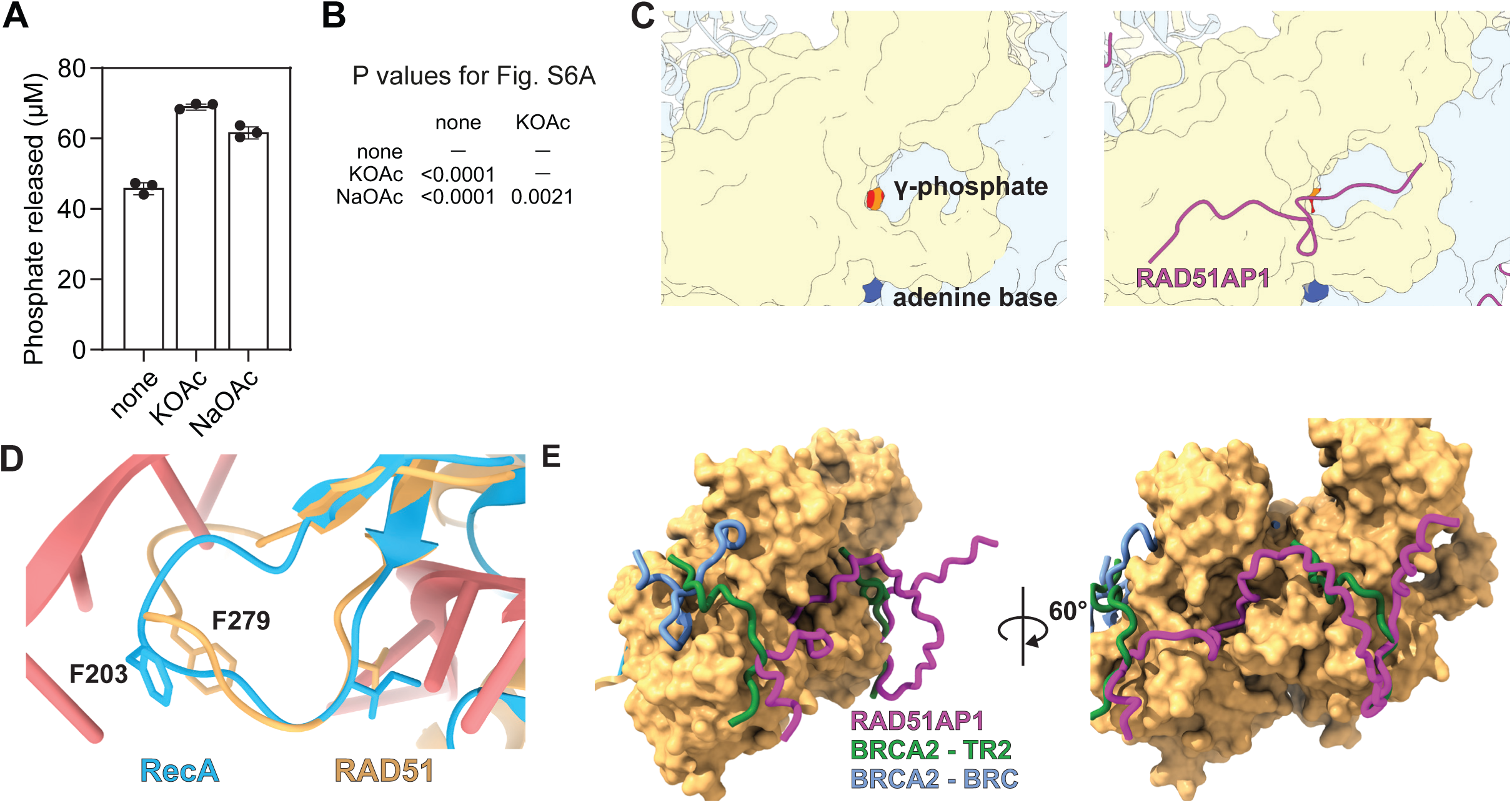
**(A)** ATPase assay showing the stimulation of RAD51 ATPase activity by 50 mM salt. potassium acetate (KOAc). sodium acetate (NaOAc). **(B)** P values for results in Figure S6A. Statistical test was one-way ANOVA followed by Tukey’s correction for multiple comparisons. **(C)** View of a possible release channel for the γ-phosphate in the Mg^2+^-ATP filament (left) and the Mg^2+^-ATP filament bound by RAD51AP1-C29 (right). **(D)** View of the L2 loop in an overlay of the structure of the RecA filament (PDB: 7iy7) during homologous recombination and the structure of the RAD51 filament bound by RAD51AP1-C29. **(E)** Overylay of the structures of RAD51 filament bound by BRCA2 TR2 (8pbc), RAD51AP1-C59 (AF3 prediction as in Figure 2I) and BRCA2 BRC4 (1n0w).

**Table S1.** Filament helical parameters of various RAD51-ssDNA filaments.

| protein | DNA | Metal ion | nucleotide | Resolution (Å) | Twist (°) | Rise (Å) | Pitch (Å) | Units per turn | reference |
| --- | --- | --- | --- | --- | --- | --- | --- | --- | --- |
| RAD51 | ss | Ca<sup>2+</sup> | ATP | 3.8 | 56.7 | 16.6 | 105.4 | 6.4 | (Appleby et al., 2023a) |
| RAD51 | ss (no density observed) | Ca<sup>2+</sup> | ADP | 3.6 | 53.1 | 18.6 | 126 | 6.8 | (Appleby et al., 2023a) |
| RAD51-BRCA2 (TR2 domain) | ss | Ca<sup>2+</sup> | ATP | 2.93 | 56.2 | 16.1 | 103 | 6.4 | (Appleby et al., 2023b) |
| RAD51-RAD51AP1 | ss | Ca<sup>2+</sup> | ATP | 3.1 | 55.9 | 16.0 | 103 | 6.4 | this study |
| RAD51 | ss | Mg<sup>2+</sup> | ATP | 3.3 | 55.1 | 16.4 | 107 | 6.5 | this study |
|  | ss (no density observed) | Mg<sup>2+</sup> | ADP | 3.6 | 52.9 | 18.5 | 126 | 6.8 | this study |
| RAD51-RAD51AP1 (C29) | ss | Mg<sup>2+</sup> | ATP | 3.1 | 55.8 | 15.9 | 103 | 6.5 | this study |

## STAR Methods

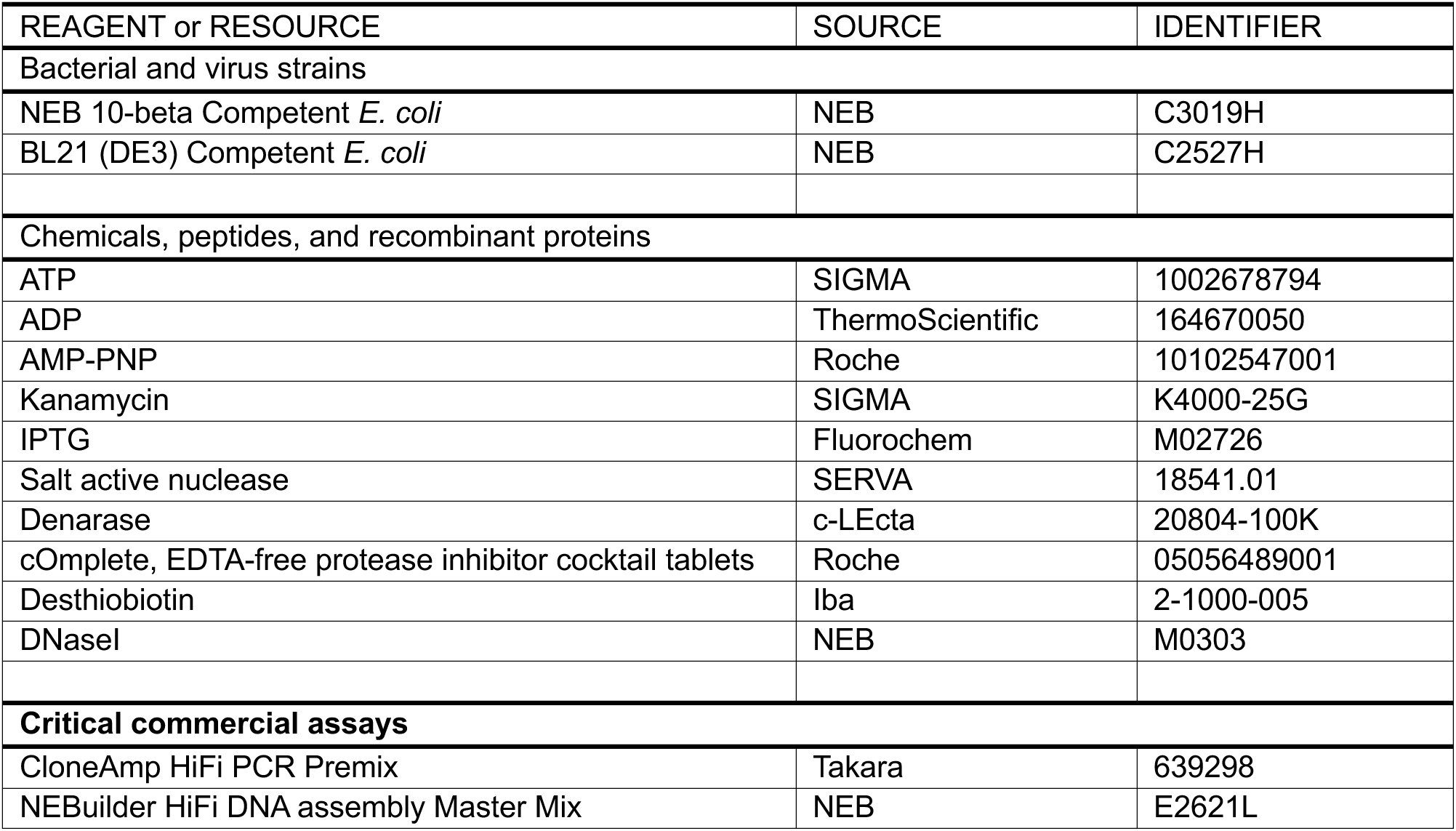
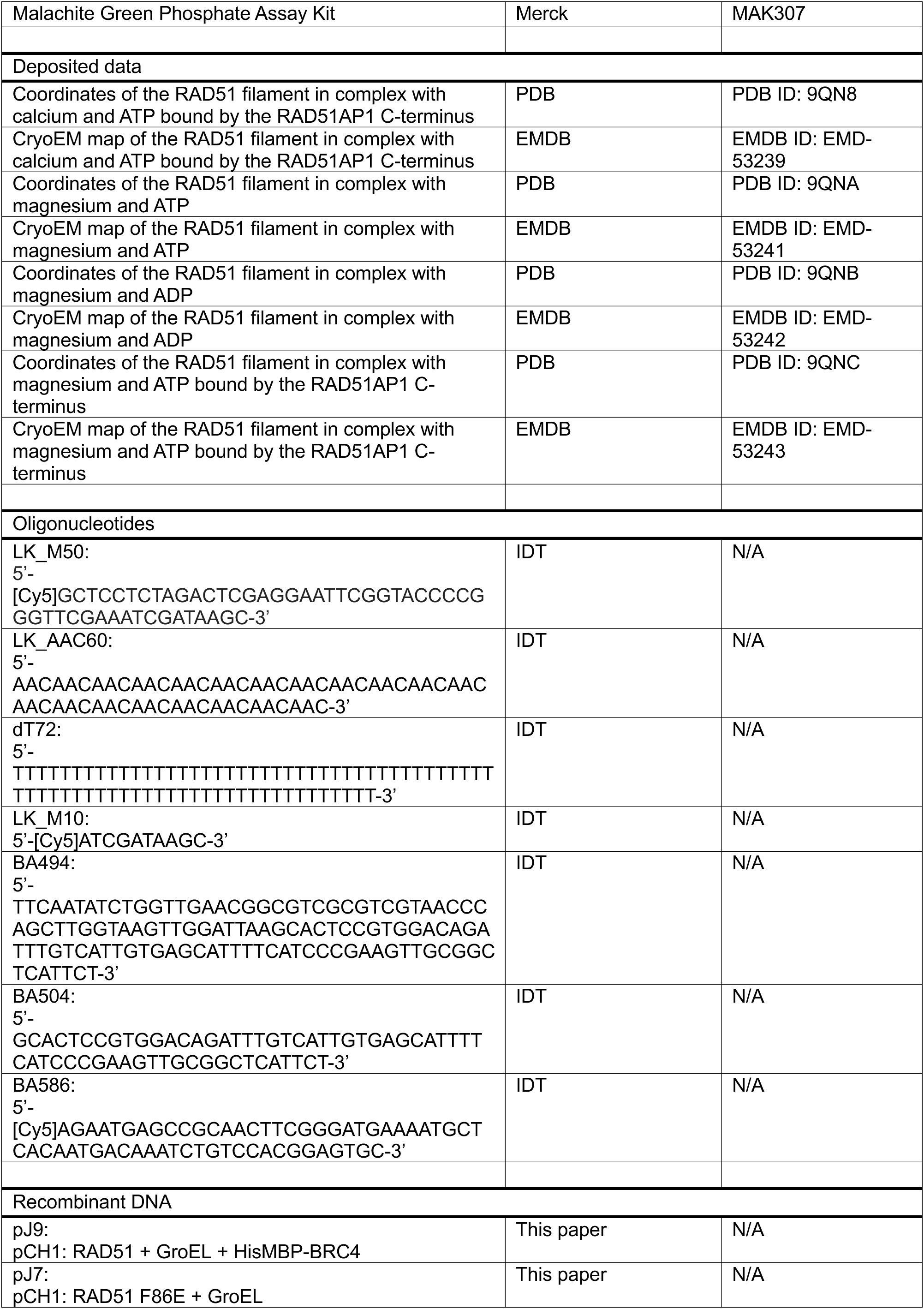
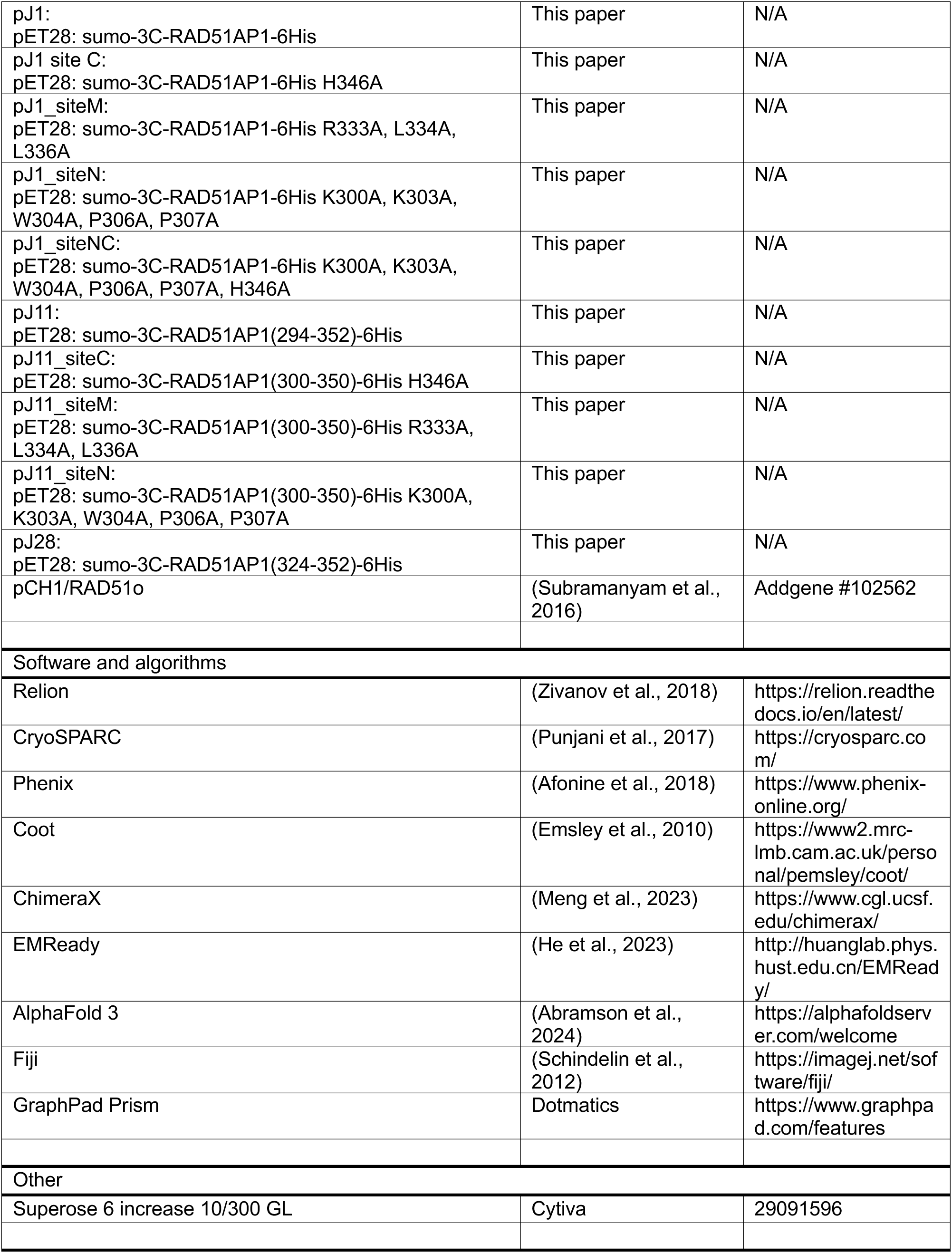

### Sequence alignment

A multiple sequence alignment of RAD51AP1 sequences of representative animals was generated using clustal omega (Sievers and Higgins, 2018) and a sequence logo was generated using the WebLogo server (Crooks et al., 2004).

### Cloning

The RAD51AP1 sequence was ordered as a synthetic gene from Invitrogen, amplified by PCR using the Cloneamp mastermix (Takara) and inserted into a pET28 vector with an N-terminal sumo tag and a C-terminal His tag. Substitution mutations and truncations were generated by PCR and the fragments were combined using the NEBuilder HiFi assembly kit (NEB).

The RAD51 F86E plasmid was generated from a pCH1 RAD51 plasmid (Subramanyam et al., 2016). The plasmid for co-expression of BRC4 with RAD51 (pJ9) was generated by amplifying 6xHis-MBP-BRC4 (Carver et al., 2024) by PCR and inserting it in the pCH1 plasmid upstream of RAD51.

### Protein purification

All proteins were expressed in BL21 (DE3) cells carrying the plasmid grown in 500 ml of LB media supplemented with kanamycin (50 μg/ml) at 37 °C to an optical density of 0.4 and then transferred to 18 °C. Protein expression was induced one hour later by addition of 0.5 mM IPTG. Cells were harvested by centrifugation and stored at -20 °C.

RAD51 was purified using an MBP-BRC4 construct as described previously with some modifications (Brouwer et al., 2018). Briefly, RAD51, His-MBP-BRC4 and GroEL were co-expressed from a single plasmid. After initial pulldown using a HisTrap column (Cytiva^TM^), RAD51 was further purified with a HiTrap Heparin column (Cytiva^TM^) and a HiTrap Q anion exchange column (Cytiva^TM^). Peak fractions were concentrated and flash frozen in liquid nitrogen.

RAD51 F86E was purified as described previously (Paoletti et al., 2020) except that the final purification was carried out using size-exclusion chromatography with an S200 increase column (Cytiva^TM^) in 50 mM Tris pH 7.5, 150 mM KCl, 1 mM EDTA, 0.5 mM TCEP.

Full length RAD51AP1 and mutants were expressed from a pET28 vector including an N-terminal sumo tag and a C-terminal his tag. Cell pellets were resuspended in His binding buffer (20 mM HEPES, pH 7.5, 500 mM KCl, 12.5 mM imidazole, 0.5 mM TCEP) supplemented with denarase nuclease (c-LEcta), salt active nuclease (SERVA) and cOmplete protease inhibitors (Roche^TM^). Cells were lysed by sonication and the lysate was clarified by centrifugation (50,000 g, 60 minutes) and then applied to a 5 ml HisTrap HP column (Cytiva^TM^). The column was washed with 20 ml His binding buffer, 10 ml wash buffer 1 (20 mM HEPES, pH 7.5, 1M KCl, 25 mM imidazole, 0.02% DDM, 0.5 mM TCEP), 20 ml wash buffer 2 (20 mM HEPES, pH 7.5, 50 mM KCl, 25 mM imidazole, 0.5 mM TCEP), 10 ml wash buffer 3 (20 mM HEPES, pH 7.5, 50 mM KCl, 300 mM imidazole, 0.5 mM TCEP) and eluted in 10 ml elution buffer (20 mM HEPES, pH 7.5, 50 mM KCl, 500 mM imidazole, 0.5 mM TCEP). The sumo tag was cleaved by adding 100 μg 3C protease and incubating at 4 °C overnight. The protein was further purified using a 1ml HiTrap Heparin column run in 20 mM HEPES, pH 7.5, 0.5 mM TCEP and KCl, eluting at a concentration of approximately 150 mM KCl. The peak fractions were pooled, diluted in the heparin running buffer and applied to a 1ml HiTrap Q column. The protein eluted at a concentration of approximately 200 mM KCl. Finally, the protein was polished by gel filtration in a superose 6 increase column (Cytiva^TM^) in 20 mM HEPES, pH 7.5, 0.5 mM TCEP and 150 mM KoAc. The protein was concentrated and frozen in aliquots in liquid nitrogen.

RAD51AP1 C59 was expressed from a pET28 vector including an N-terminal sumo tag and a C-terminal his tag. Cell pellets were resuspended in buffer A (20 mM Tris, pH 8, 50 mM NaCl, 0.5 mM TCEP) supplemented with 10 mM imidazole, denarase nuclease (c-LEcta) and cOmplete protease inhibitors (Roche^TM^). Cells were lysed by sonication and the lysate was clarified by centrifugation (75,000 g, 30 minutes) and then applied to a 5 ml HisTrap HP column (Cytiva^TM^). The column was washed with 10 ml buffer B (20 mM Tris, pH 8, 1000 mM NaCl, 0.5 mM TCEP) supplemented with 10 mM imidazole and 0.02% DDM, followed by 15 ml of buffer A supplemented with 50 mM imidazole. Proteins were eluted with buffer A containing 400 mM imidazole. The sample was further purified using a HiTrap Q column (Cytiva^TM^) and eluted in a gradient of 100 mM to 600 mM NaCl in buffer A. Finally, the protein was polished by gel filtration in a superose 6 increase column (Cytiva^TM^) in 20 mM HEPES, pH 7.5, 0.5 mM TCEP and 150 mM KoAc. The protein was concentrated and frozen in aliquots in liquid nitrogen.

RAD51AP1 C29 was expressed from a pET28 vector with an N-terminal twin strep-sumo tag and a C-terminal His tag. Cell pellets were resuspended in buffer TBK300 (20 mM Tris, pH 7.5, 300 mM KCl, 0.5 mM TCEP) supplemented with denarase nuclease (c-LEcta), salt active nuclease (SERVA) and cOmplete protease inhibitors (Roche^TM^). Cells were lysed by sonication and the lysate was clarified by centrifugation (50,000 g, 30 minutes) and then applied to a 5 ml StrepTrap HP column (Cytiva^TM^). The column was washed with 10 ml wash buffer (20 mM Tris, pH 7.5, 1000 mM KCl, 0.5 mM TCEP), 10 ml TBK300 and eluted in 15 ml TBK300 containing 2 mg/ml desthiobiotin. The twin strep and sumo tag were removed using 100 μg 3C protease, which were added to the eluate and incubated overnight at 4 °C. The sample was then diluted to 100 mM KCl and applied to a HiTrap Q column (Cytiva^TM^). The protein was found in the flow-through, which was concentrated and frozen in liquid nitrogen.

### Pull-down assays

Hexahistidine-tagged RAD51AP1 (1 µM of full-length or 2 µM of C59) in 30 µL of pull-down buffer (25 mM Tris-OAc [7.5], 150 mM or 100 mM KOAc [full-length or C59, respectively], 1 mM ATP, 5 mM MgOAc, 0.25 mM TCEP, 0.1 mg/mL BSA, 0.1% Tween 20, 2.5% glycerol, 50 mM imidazole) was immobilized on magnetic cobalt resin (ThermoFisher Scientific^TM^ 10103D) by incubating at 37°C for 10 min with mixing. A magnetic stand was used to discard the supernatant. The protein-bound resin was then resuspended in 30 µL of pull-down buffer containing RAD51 (1 µM for full-length and 2 µM for C59) and incubated with mixing at 37°C for 5 min. The supernatant containing unbound RAD51 was recovered and bound proteins were eluted by incubation with SDS-PAGE loading buffer (65°C, 1300 rpm, 5 min). The protein content of both fractions was analysed by SDS-PAGE and staining (ReadyBlue Sigma-Aldrich), and gels were imaged using a BioRad imager (ChemiDoc MP Imaging System^TM^). FIJI was used for quantification (Schindelin et al., 2012). Briefly, the images were background subtracted using the rolling ball method (50 pixels), then the intensity of RAD51 and RAD51AP1 bands in the supernatant and eluate was determined. The percentage of signal in the eluate was calculated. For each experiment, the value for the non-specific RAD51 binding (minus RAD51AP1 sample) was subtracted from all samples, then the amount of RAD51 was normalized to RAD51AP1 binding and expressed relative to wild-type protein.

### Bio-layer interferometry (BLI)

Bio-Layer Interferometry (BLI) experiments were performed on an Octet Red instrument (Sartorius) operating at 25 °C. RAD51AP1 (full-length or C59, wild-type or mutant variants) with a C-terminal hexahistidine tag was diluted to 1 µg/mL with BLI buffer (25 mM HEPES [7.5], 150 mM KOAc, 5 mM MgOAc, 1 mM ATP, 0.05% Tween20, 0.5 mM TCEP) and transferred into a row on a 96 well plate (200 µL per well, x8). RAD51 was serially diluted (two-fold) with BLI buffer from 2.5 µM to 39 nM, and along with a buffer alone sample, 200 µL of each concentration was transferred into a separate row. Nickel-NTA biosensors (Sartorius cat. # 18-5101) were soaked in BLI buffer for 20 min prior to experiments. The following steps were automated. First, the biosensors were dipped in BLI buffer (100 s) for a first Baseline step, then RAD51AP1 was loaded onto the biosensors until a response of 0.1 nm was achieved. Next, the biosensors were transferred into fresh BLI buffer for a second Baseline (40 s), to check RAD51AP1 was stably bound to the sensors. The biosensors were then transferred into the RAD51 dilution series to monitor association and then to fresh BLI buffer to monitor dissociation. Double reference subtraction was performed for all samples by subtracting the curve recorded without RAD51 and the response values recorded with no RAD51AP1 loaded on the sensors. The resultant responses were averaged and then plotted against RAD51 concentration. Data were analysed using Octet BLI Analysis software (Sartorius) and in-house software (Martin et al., 2021). Equilibrium dissociation constants (Kd) were estimated from the instrument response against RAD51 concentration using least squares non-linear regression and a 1:1 binding model. Experiments were repeated in triplicate and errors are reported as standard deviations from the mean.

### Nuclease protection assay

10 µMnt of a Cy5-labeled oligonucleotide (60-mer, BA586) in nuclease protection buffer (25 mM Tris-OAc [7.5], 100 mM KOAc, 2 mM ATP, 5 mM MgOAc, 0.5 mM TCEP, 2.5% glycerol) was supplemented with RAD51 (2 µM) and the indicated concentration of RAD51AP1. Following a 5 min incubation at 37°C, 2 µL of DNase I (NEB^TM^ M0303) was added and incubation continued for 15 min. Reactions were then deproteinized by addition of stop solution (final concentration 12 mM Tris-Cl [7.5], 7.52 mM EDTA, 0.45% SDS, 0.75 mg/mL proteinase K). DNA was resolved on 10% PAGE gels and imaged using a Bio-Rad imager (ChemiDoc MP Imaging System^TM^). FIJI was used for quantification (Schindelin et al., 2012). Briefly, the images were background subtracted using the rolling ball method (50 pixels). Next, the total lane signal was determined and the fraction of signal corresponding to intact DNA was then expressed as a percentage of the total.

### Negative stain

Filaments were formed on oligo BA494 (1.5 μM nucleotides) with RAD51 (375 nM) and C59 (1.5 μM) in buffer D (20 mM HEPES, pH 7.5, 100 mM KCl, 0.5 mM TCEP) supplemented with 2 mM ATP and 5 mM MgCl2 at 37 °C for 5 minutes. 10 μl samples were applied to glow discharged 300 mesh carbon coated copper grids (Agar Scientific), blotted and stained with 2% uranyl acetate. Micrographs were acquired on a T12 microscope (FEI) at a magnification of 21,000 using a Rio CMOS camera (Gatan^TM^). Filament lengths were measured as described previously (Greenhough et al., 2023).

### Strand exchange assay

RAD51 (1 µM) was incubated with 3 µMnt of a 116-mer oligonucleotide (BA494) in strand exchange buffer (25 mM Tris-OAc [7.5], 100 mM KOAc, 2 mM ATP, 1 mM MgCl2, 2 mM CaCl2, 0.5 mM TCEP, 0.1 mg/mL BSA) at 37°C for 5 min in the presence of the indicated concentration of RAD51AP1. RPA (0.2 µM) was then added to the reaction, and following another incubation (37°C, 5 min), the reaction was intitated via the addition of 1 µMbp 60-mer dsDNA (BA586-BA504) and incubated at 37°C for 30 min. Reactions were deproteinized, resolved by PAGE, and imaged as above. FIJI was used for quantification (Schindelin et al., 2012). Briefly, the images were background subtracted using the rolling ball method (50 pixels). Next, the total lane signal was determined and the fraction of signal corresponding to product was then expressed as a percentage of the total. This was then expressed relative to the RAD51 alone reaction to yield fold stimulation.

### ATPase assay

3 µM RAD51 and 9 µMnt poly-dT ssDNA (72-mer) were incubated in ATPase buffer (25 mM Tris-OAc [pH 7.5], 0.5 mM ATP, 5 mM MgOAc, 0.5 mM TCEP, 0.01% Tween20, 2.5% glycerol), supplemented with 50 mM salt (KOAc or NaOAc, as indicated), at 37°C for 90 min. Reactions were stopped by addition of 20 mM EDTA then diluted four-fold to reduce phosphate concentration. A commercial colorimetric kit was then used following the manufacturer’s instructions (MAK307, Merck) to determine phosphate concentration.

### Electrophoretic mobility assay

1 µM LK_M10 in buffer D containing AMP-PNP (2 mM) and MgCl2 (5 mM) was mixed with RAD51 (4 µM) and C29 (10 µM) or buffer as indicated. After incubation for 5 minutes at room temperature the mixture was resolved by native PAGE.

### cryoEM sample preparation

#### RAD51AP1 bound to RAD51 filament formed with Ca^2+^-ATP

Filaments were formed on oligo LK_M50 (12 μM nucleotides) with RAD51 (4 μM) in buffer D supplemented with 2 mM ATP and 5 mM CaCl2 at 37 °C for 5 minutes. Immediately before preparing grids, the filaments were diluted to a final concentration of 1.6 μM RAD51. 2 μl of filaments were applied to glow discharged 300 mesh carbon coated lacey carbon grids (LC300-Au-UL, EM Resolutions) in a Mark IV vitrobot (ThermoFisher Scientific^TM^) and incubated at 24 °C, 100% humidity for 3 minutes. 1 μl of sample was removed from the grid and 2 μl of RAD51AP1 (0.8 μM in buffer D supplemented with 2 mM ATP and 5 mM CaCl2) was applied to the grid and incubated for 1 min. Finally, 2 μl were removed and replaced with 3 μl of buffer and incubated for 1 minute. Grids were then blotted and plunged into liquid ethane.

#### RAD51 filament formed with Mg^2+^-ATP

Filaments were formed on oligo LK_AAC60 (21 μM nucleotides) with RAD51 (10.5 μM) in buffer D supplemented with 3 mM ATP and 7.5 mM MgCl2 at 37 °C for 5 minutes. They were then diluted 2:1 with buffer D containing ATP and MgCl2 to a final RAD51 concentration of 7 μM. 4 μl of filaments were applied to R2/2, 200 mesh ultrafoil grids (EMS^TM^), which had been washed in chloroform before use, in a Mark IV vitrobot (ThermoFisher Scientific^TM^) at 4 °C, 100% humidity and blotted and plunged into liquid ethane after 30 seconds.

#### RAD51AP1 C29 bound to RAD51 filament formed with Mg^2+^-ATP

The filaments were formed as above and then diluted 2:1 with C29 in buffer D containing ATP and MgCl2 to final RAD51 concentration of 7 μM and a final concentration of C29 of 28 μM. The grids were then prepared in the same manner as for the apo filament.

All grids were imaged at the LonCEM facility at the Francis Crick Institute on a Titan Krios microscope equipped with a K3 detector (Gatan^TM^) at a pixel size of 1.08 Å. Full acquisition details are in **Table 1**.

### cryoEM data processing

For all datasets, movies were processed with MotionCor2 as implemented in relion5 (Zivanov et al., 2019). All subsequent processing was done in cryoSPARC version 3-4.6 (Punjani et al., 2017).

#### RAD51AP1 bound to RAD51 filament formed with Ca^2+^-ATP

5001 micrographs were imported into cryoSPARC. CTF was estimated using PatchCTF and 159 micrographs were rejected based on poor CTF fit resolution worse than 10 Å and defocus lower than 0.3 μm. Particles were picked using template free filament tracer. 2.4 million particles were picked and extracted binned 4 times. After multiple rounds of 2D classification in cryoSPARC, 510,927 particles were selected and re-extracted binned 2 times. After further 2D classification, 489,416 particles were selected and re-extracted without binning. After further 2D classification, 458,792 particles were selected. Homogeneous 3D refinement, without applying helical symmetry, using an initial volume of an AP1 bound filament from a previous screening dataset led to a map of the filament at a resolution of 3.1 Å. A mask was designed around a piece of weak density on the outside of the filament. After three rounds of 3D classification with 10 classes each followed by homogeneous refinement, 137,283 particles were selected and refined with homogeneous refinement, resulting in a map at a resolution of 3.2 Å. Further minimizing per-particle scale and per-particle defocus improved resolution to 3.1 Å. The interpretability of the map was further improved using EMReady (He et al., 2023). Final helical parameters were estimated based on the central part of the map in which the atomic model was built, giving rise to final estimates of a helical rise of 16 Å and a helical twist of 55.9°.

#### RAD51 filaments formed with Mg^2+^-ATP

3096 micrographs were imported into cryoSPARC. CTF was estimated using PatchCTF and 224 micrographs were rejected based on poor CTF fit resolution worse than 10 Å and defocus lower than 0.3 μm. 6.6 million particles were picked using a combination of template free filament tracer and template picker and template based filament tracer using 2D averages of a RAD51 filament from a previous screening dataset as templates. After removing overlapping particles, 5.6 million particles were extracted in a small box (180 pixels) and after 2 rounds of 2D classification 983,859 particles were selected and re-extracted in a larger box (256 pixels), binned 2 times. After further 2D classification, 538,943 particles were selected and refined with helical refinement, using a reconstruction of the RAD51AP1 bound Ca^2+^-ATP RAD51 filament as an initial volume and initial helical twist estimate of 56° and a rise estimate of 16 Å. Refinement of the helical parameters during helical refinement led to revised estimates of 54.6° and 16.9 Å respectively. These were used as input values into another helical refinement, resulting in final estimates of 54.1° and 17.3 Å in the consensus refinement. The particles were classified in 3D into 5 classes and the particles re-extracted without binning and then subjected to helical refinement again. The 3D classes were grouped into two groups, one with high helical rise and one with low rise and particles belonging to each group were combined and refined again. They were then classified again in 2D and 229,610 particles belonging to the high helical rise conformation and 242,944 particles belonging to the low helical rise conformation were selected. Final helical refinement resulted in volumes at resolutions of 3.6 Å and 3.3 Å respectively. The interpretability of the maps was further improved using EMReady (He et al., 2023). Final helical parameters were estimated based on the central part of the map in which the atomic model was built, giving rise to final estimates of a helical rise of 18.5 Å and a helical twist of 52.9° and a helical rise of 16.4 Å and a helical twist of 55.1°.

#### RAD51AP1 C59 bound to RAD51 filament formed with Mg^2+^-ATP

3233 micrographs were imported into cryoSPARC. CTF was estimated using PatchCTF and 812 micrographs were rejected based on poor CTF fit resolution worse than 10 Å and defocus lower than 0.3 μm. 7.5 million particles were picked using a combination of template free filament tracer and template picker and template based filament tracer using 2D averages of a RAD51 filament from a previous screening dataset as templates. After removing overlapping particles, 6.1 million particles were extracted in a small box (180 pixels) and after 2 rounds of 2D classification 1.9 million particles were selected and re-extracted in a larger box (256 pixels), binned 2 times. After further 2D classification, 1.2 million particles were selected and refined with helical refinement, using a reconstruction of the RAD51AP1 bound Ca^2+^-ATP RAD51 filament as an initial volume and initial helical twist estimate of 56° and a rise estimate of 16 Å. Refinement of the helical parameters during helical refinement led to revised estimates of 55.3° and 16.5 Å respectively. The particles were classified in 3D with a small mask around the density outside of the filament on the central protomer. Two classes with the clearest additional density were combined, and the particles were re-extracted without binning, refined using homogeneous refinement, local CTF refined, and classified in 2D again. The resulting volume was again subjected to 3D classification with a mask around the central RAD51 protomer. Each class was refined using homogeneous refinement and the highest resolution class was selected and classified in 2D again, resulting 97,029 particles. Homogeneous refinement with minimizing per-particle scale and per-particle defocus resulted in a final volume at a resolution of 3 Å. Final helical parameters were estimated based on the central part of the map in which the atomic model was built, giving rise to final estimates of a helical rise of 15.9 Å and a helical twist of 55.8°.

### Model building

Models of 6 RAD51 protomers with ssDNA, ATP or ADP, Ca2+ or Mg2+ and RAD51AP1 C29 as appropriate were modelled in AlphaFold (Abramson et al., 2024) and docked into the maps as rigid bodies. The models were partially manually rebuilt in Coot (Emsley et al., 2010). The final models were refined using phenix.real_space_refine (Afonine et al., 2018).

### Estimation of helical parameters

Each map was split to extract the central part in which the model had been built using ChimeraX (Meng et al., 2023). This map was then imported into cryoSPARC and the helical parameters were estimated using the job symmetry search utility.

## Notes

### Competing Interest Statement

The authors have declared no competing interest.

